# Living Cells Employ Ubiquitin-Proteasomal System and Nucleotide Excision Repair Pathways to Remove Reactive Oxygen Species-Induced DNA-Protein Crosslinks (ROS-DPCs)

**DOI:** 10.64898/2026.02.06.704426

**Authors:** Cesar I Cyuzuzo, Monica Kruk, Qi Zhang, Duha Alshareef, James Harmon, Yuichi J Machida, Harrison W. VanKoten, Swati S More, Colin Campbell, Natalia Y Tretyakova

## Abstract

Oxidative DNA damage caused by endogenous reactive oxygen species (ROS) is a key driver of mutagenesis, cellular dysfunction, and aging, contributing to diseases like cancer, neurodegeneration, rheumatoid arthritis, cardiovascular disorders, and diabetes. Although more than 20 oxidative base lesions have been identified, ROS-induced DNA-protein crosslinks (DPCs) are poorly characterized. ROS-DPCs are unusually bulky and highly toxic lesions that accumulate in metabolically active tissues with age, but their identities, biological consequences, and repair in living cells have remained elusive. In the present work, we characterized ROS-DPCs in human fibrosarcoma (HT1080) cells treated with hydrogen peroxide (H_2_O_2_) and elucidated the mechanisms of their removal. Mass spectrometry-based proteomics has identified over 100 cellular proteins that participated in DPC formation, most of which are involved in DNA metabolism. Our data further reveal that DNA replication and transcription facilitate DPC detection and identify a critical role of the ubiquitin-proteasomal system (UPS), replication-coupled activity of SPRTN metalloprotease, and nucleotide excision repair (NER) in removing ROS-induced DPCs. ROS-DPC formation was blocked by pretreatment with metabolically stable and cell-permeable glutathione (GSH) analog (Ψ-GSH), suggesting a possible therapeutic strategy for preventing diseases associated with increased ROS levels.

**KEY POINTS:** Mass spectrometry-based proteomics identified over 100 proteins participating in DNA-protein cross-links in human cells treated with ROS

Our work reveals the mechanisms through which living cells recognize and remove ROS-DPCs

Our study demonstrates the potential of a glutathione analog to prevent ROS-DPC formation

**GRAPHICAL ABSTRACT:** 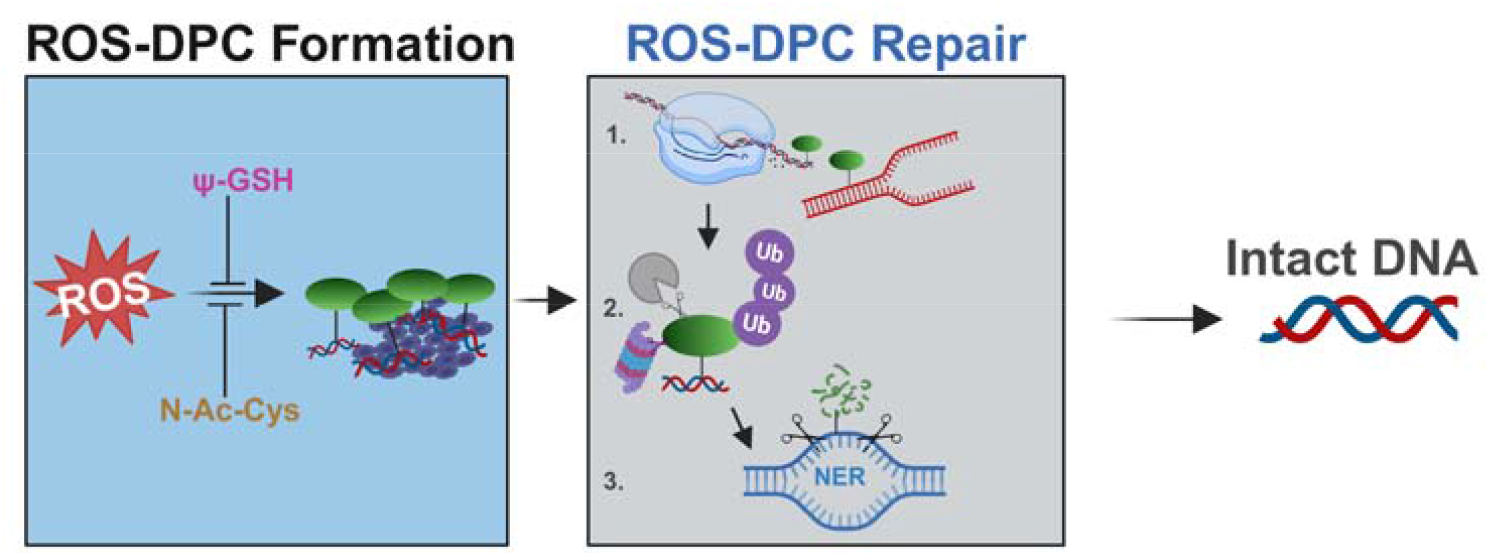

## INTRODUCTION

DNA-protein crosslinks (DPCs) are unusually bulky, ubiquitous DNA lesions that arise when cellular proteins become covalently trapped on DNA. DPCs can be induced upon exposure to endogenous or exogenous cross-linking agents and can also form as a result of enzymatic trapping of topoisomerases, DNA polymerases, and DNA repair proteins on DNA strands (1–3).

Common DPC-inducing agents include endogenous cellular metabolites such as formaldehyde (4), methylglyoxal (5), malondialdehyde, and reactive oxygen species (ROS) (6,7). Given their considerable size and their ability to disrupt normal DNA-protein interactions, unrepaired DPCs obstruct nearly all DNA transactions, including repair, transcription, and replication, ultimately leading to cell death. DPCs have been shown to contribute to the antitumor activity of many anticancer agents, including platinum agents and nitrogen mustards (8,9).

Endogenously formed DPCs have been implicated in aging, neurodegenerative conditions, and cardiovascular disease (10–12). However, the mechanisms of formation and identities of the participating proteins remain elusive. DPC levels are reported to increase significantly with age (10), probably due to endogenous exposure to reactive oxygen species (ROS) and lipid peroxidation products (7,13). Mice deficient in SPRTN, a specialized metalloprotease involved in DPC processing, are not viable, while germline mutations in SPRTN result in Ruijs-Aalfs syndrome (RJALS) in humans, a disorder characterized by genome instability, progeria, and early-onset hepatocellular carcinoma (14,15). Several mechanisms could be involved in endogenous DPC formation, including enzymatic trapping of topoisomerases, polymerases, and repair proteins on DNA (16,17), exposure to endogenous *bis*-electrophiles such as methylglyoxal and formaldehyde (5,18), and free-radical mechanisms involving endogenous ROS (19).

Examples of physiologically relevant ROS include superoxide anions, hydroxyl radicals, carbonate radical anions, nitrosoperoxycarbonate, and hydrogen peroxide (H_2_O_2_) (20,21). While H_2_O_2_ is poorly reactive towards DNA, it can be activated by reactions with iron (II) to produce other potent DNA-damaging ROS, such as carbonate radical anion and hydroxyl radical (22). One common type of DPC lesions formed upon exposure to ROS involves the thymine residues of DNA and the tyrosine side chains of proteins (2’-deoxythymidine-tyrosine (dT-Tyr) (**Scheme 1**)). dT-Tyr has been used as a biomarker of ROS-DPC in several cell and tissue models (19,23). Our previous study demonstrated that genomic levels of dT-Tyr DPCs were increased in cardiomyocytes of rats that underwent ligation/reperfusion surgery as a model of ischemia-reperfusion injury following a heart attack (24). Mass spectrometry (MS) based proteomics revealed that nearly 90 proteins participated in DPC formation in the heart following reperfusion injury (24). Major classes of crosslinked proteins included ROS-scavenging enzymes (e.g., catalase and superoxide dismutase), sarcomere proteins (e.g., myosin-6, 7, and 9), and other abundant cellular proteins (e.g., actin, histones) (24,25). Interestingly, approximately 43% of oxidatively cross-linked proteins were of mitochondrial origin, highlighting the increased production of superoxide anions, H_2_O_2_, and hydroxyl radicals in this organelle (20,26). We have also quantified dT-Tyr formed in murine tissues endogenously and following exposure to ionizing radiation (7), but the identities of the participating proteins have not been elucidated.

Given inherent structural heterogeneity of DPC adducts and their deleterious cellular effects, an integrated cellular response to these lesions is required. DPC repair requires ubiquitin ligases, DPC proteases, nucleases, and DNA damage recognition proteins (27–31). Via a direct reversal mechanism, tyrosyl-DNA phosphodiesterases (TDP) 1 and 2 can hydrolyze the covalent linkage between DNA and topoisomerases 1 and 2, respectively (29,32-34). However, the majority of DPCs formed in human cells cannot be reversed and require proteolytic digestion of DNA-crosslinked proteins, which can be accomplished by SPRTN metalloprotease or the proteasome (14,35-39). We recently reported that mice deficient in SPRTN metalloprotease accumulate dT-Tyr lesions (7). Similarly, human cells deficient in SPRTN were sensitized to ROS and contained higher levels of dT-Tyr (15). Following proteolytic processing of DPCs, the remaining DNA-peptide lesions can be removed via canonical DNA repair pathways, such as nucleotide excision repair (NER) (18,40,41), or tolerated by homologous recombination (HR) or the translesion synthesis pathway (42,43). Additionally, several other enzymes, including nucleases (e.g., MRE11, CtIP, Apex 1 and 2 (44,45), proteases (e.g., FAM11A (46), ACRC/GCNA (46), DDI1 and 2 (47,48)), and ATPases (e.g., p97 and its cofactor, TEX264 (38)), have been implicated in the processing of cellular DPCs.

While SPRTN protease detects DPCs during DNA replication and binds to ssDNA-dsDNA junctions, such as those found at stalled replication forks (36,49), non-replicating cells such as neurons require alternative mechanisms for DPC processing. Emerging evidence supports the role of the ubiquitin-proteasomal system (UPS) in DPC removal (28,36,48,50). *Sun et al*. reported that topoisomerase-induced DPCs accumulate in mammalian cells co-treated with topoisomerase poisons and ubiquitin ligase inhibitor (28). Additionally, Larsen et al. demonstrated that ubiquitination facilitates the recruitment of SPRTN metalloprotease and the proteasome to DNA methyltransferase-DPCs in Xenopus egg extracts (36). Yet, less is known about non-enzymatic ROS-DPCs repair in human cells and the respective roles of individual repair factors.

The goal of the present study was to characterize ROS-mediated DNA-protein crosslinking in human cells treated with hydrogen peroxide and to elucidate the mechanisms of their processing and removal. We employed cell models deficient in the NER pathway and leveraged pharmacological inhibition of DPC recognition factors to establish the relative contributions of these mechanisms in ROS-DPC repair. MS-based proteomics enabled the identification of the proteins that participate in ROS-DPC formation. Our *in vitro* experiments corroborated the formation of ROS-DPC by high-mobility group B2 protein (HMGB2) in the presence of H_2_O_2_ and a DNA duplex. Finally, we tested the ability of a metabolically resilient synthetic glutathione analog (ψ-GSH) (51,52) to prevent DPC formation in human cells under oxidative stress conditions. Collectively, our results could contribute to the understanding of the molecular mechanisms underlying the cellular response to ROS-DPCs in pathologies associated with oxidative stress, including aging, cardiovascular disease, cancer, neurodegenerative diseases, diabetes, and immune dysfunction.

**Scheme 1:**
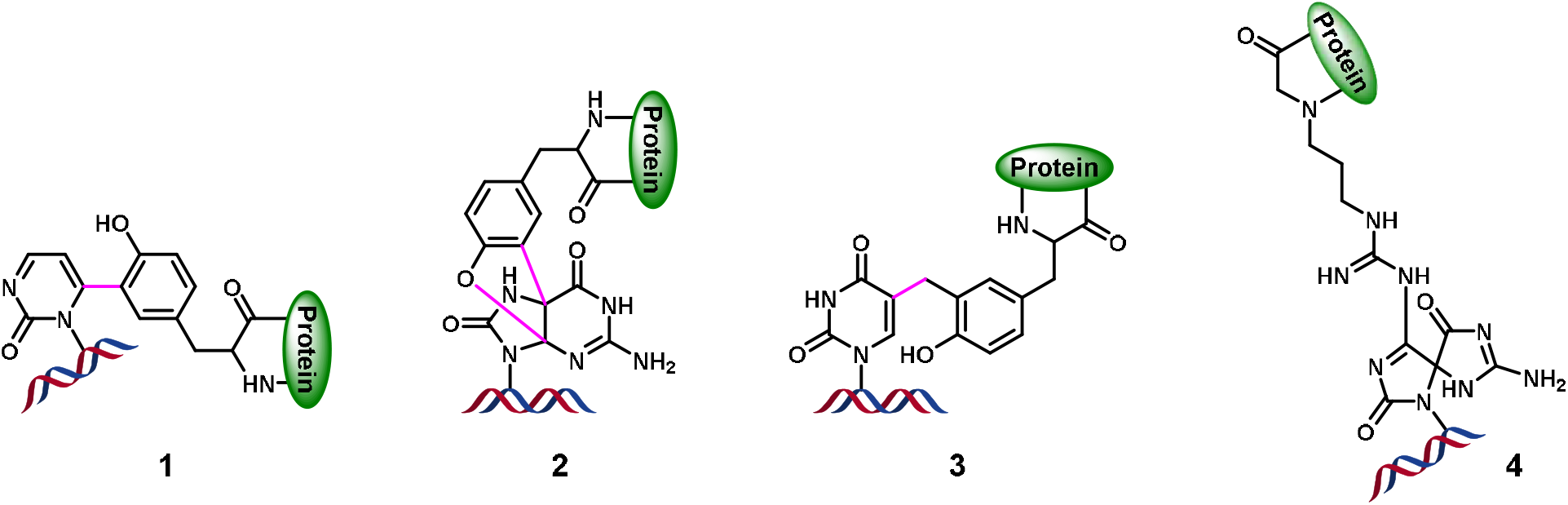
Chemical structures of DNA-protein cross-links induced by reactive oxygen species (ROS). **1: 6**-(2-hydroxy-5-(2-(methylamino)-3-oxobutyl)phenyl)-1-methylpyrimidin-2(1H)-one, **2:** 2-amino-3-(2-amino-10-(4-hydroxy-5-(hydroxymethyl)tetrahydrofuran-2-yl)-4,11-dioxo-3,4-dihydro-4a,9a (epiminomethanoimino)benzofuro[2,3-d]pyrimidin-6-yl) propanoic acid, **3:** 2-amino-3-(4-hydroxy-3-((1-((2R,4R,5R)-4-hydroxy-5-(hydroxymethy)tetrahydrofuran-2-yl)-2,4-dioxo-1,2,3,4-tetrahydropyrimidin-5-yl) methyl) propanoic acid **(dT-Tyr). 4:** N6-(5-guanidino-1-(4-hydroxy-5-(hydroxymethyl) tetrahydrofuran-2-yl)-2-oxo-2,3-dihydro-1H-imidazol-4-yl) lysine.

## MATERIALS & METHODS

### Safety Statement

Phenol and chloroform are toxic solvents that should be handled with caution in a well-ventilated fume hood, wearing appropriate personal protective equipment (PPE).

### Reagents and materials

Hydrogen peroxide (H_2_O_2_ 30 wt.% in H_2_O), L-ascorbic acid (Vit C), N-acetyl-L-cysteine (N-Ac), dithiothreitol (DTT), nuclease P1 (from *penicillium citrinum*), phenol, chloroform, iodoacetamide, alkaline phosphatase, EDTA-free protease inhibitor cocktail tablets, and nuclease P1 were obtained from Sigma-Aldrich (St. Louis, MO). FeSO_4_ was purchased from Fischer Chemicals (Zurich, Switzerland). Ubiquitin-activating enzyme (UAE) inhibitor TAK-243 and 26S proteasomal subunit inhibitor, MG-132, were purchased from Fisher Scientific (Buchs, Switzerland) and Abcam (Cat. #ab14707), respectively. QuantiFluor dsDNA system kit was purchased from Promega (Madison, WI). Acetonitrile, glacial acetic acid, and hydrochloric acid were acquired from Fischer Scientific (Buchs, Switzerland). SimplyBlue™ SafeStain was bought from Invitrogen. RNase A and cell lysis solution were purchased from Qiagen (Hilden, Germany). Gibco-branded Fetal bovine serum (FBS), Dulbecco’s modified Eagle medium (DMEM), and Alpha-modified Eagle medium (α-MEM) were purchased from Thermo Fischer Scientific (Green Island, NY). Epidermal growth factor (EGF) and Fibroblast growth factor (FGF) were obtained from R&D Systems (Minneapolis, MN). Non-essential amino acids (NEAA) were purchased from Stem Cell Technologies (Cambridge, MA). Mass spectrometry grade Pierce™ Trypsin/Lys-C protease, Pierce™ BCA protein assay kit, NuPAGE 4-12% bis-tris SDS-PAGE gels, and cell culture media/reagents were obtained from Thermo Fischer Scientific (Waltham, MA). The recombinant H3.1 and proteinase K were purchased from New England Biolabs (Beverly, MA). Omega Nanosep 10K filters were acquired from PALL Life Science (Port Washington, NY), while Amicon 3K filters were obtained from Millipore (Darmstadt, Germany).

### Cell lines

Human fibrosarcoma (HT1080) cells (53) were purchased from the American Type Culture Collection (ATCC; CCL-121, https://www.atcc.org/). XPD-deficient (GM08207) cells were obtained from the Coriell Institute of Medical Research (CIMR). The HT1080-XPA-KO and HT1080-CSB-KO cells (purchased from Synthego as knockout cell pools) were obtained in the Dr. Colin Campbell’s laboratory (18). The complete loss of XPA in the HT1080-XPA-KO cell line was confirmed via western blotting **(Figure S4)**. Wildtype Sprtn^+/+^ and SPRTN-deficient (Sprtn^f/-^) MEFs were engineered as previously described (54). The cell lines above were maintained in Dulbecco’s modified Eagle medium (DMEM) supplemented with 10% FBS and 1% penicillin/streptomycin antibiotics. HT1080-XPF-KO (XP51RO cell line with a null XPF mutation) was a generous gift from Dr. Laura Niedernhofer (Univ. of Minnesota) (55) and was grown in αMEM media with 10% FBS, 1% penicillin/streptomycin, 0.002% EGF, 0.0005% FGF, and 1% NEAA. All cells were maintained in a humidified incubator at 37 °C with 5% CO_2_ and grown until a confluency of ~80% or higher was achieved. Splitting, treatments, or downstream experiments were executed within a third to eighth passage range.

### *In vitro* characterization of DPCs generated with recombinant HMGB2

Biotinylated 69-mer double-stranded DNA featuring the sequence: Biotin-5’-TTG GTA AAA CGA CGG CCA GTG CCA AGC CAT GCC TGC AGG TCG ACT CTA GAG GAT CCC CCA GCC TGC CGG-3’ (IDT Inc., 250 pmol) was incubated with a 2-fold molar excess of HMGB2 (MedChemExpess, 500 pmol) in the presence of 0, 0.1, 1, and 5 mM H_2_O_2_ in 100 mM NH_4_CO_3_ (pH 8) for 3 h at 37 °C (total reaction volume: 30 μL). DPC samples were purified using streptavidin beads and NINTA gravity columns before being visualized on 4-12% Bis-Tris SDS-PAGE Gels (Thermo Fisher Scientific; Waltham, MA).

### Alamar Blue Cell Viability Assay for H_2_O_2_ Cytotoxicity

After seeding 30,000 HT1080 cells (WT or NER KO) into 96-well plates for 24 h, the media were replaced with H_2_O_2_-containing FBS-free media for 1 h. Following treatment, the media were replaced with FBS-free DMEM, and cells were allowed to recover for 17 h. After the recovery period, the media was removed and replaced with 100 μL of a 10% alamar blue reagent dissolved in FBS-free DMEM. The 96-well plates were further incubated for 4 hours in the dark at 37 °C, and fluorescence was measured at Ex/Em 560/590 nm using a Bio-Tek Synergy H1 plate reader. The fluorescence signal for each treatment condition was normalized to the vehicle-treated control to calculate cell viability.

### Quantification of DPCs using the K-SDS Assay

DPC-containing DNA was isolated via the modified K-SDS assay (7). Briefly, cells were lysed with 0.5 mL of Buffer A [20 mM Tris-HCl (pH 7.4), 2% SDS]. Cell lysates were solubilized using 22G syringes. Lysates were treated with 200 μg/mL RNase for 15 min at 37 °C. 50 μL aliquots of the lysates were set apart as “Input DNA fractions”. The remaining 450 μL were designated as “DPC pellet fractions”. An equal volume of Buffer B [200 mM KCl in 20 mM Tris-HCl (pH 7.5)] was added to the “DPC pellet fractions” to precipitate DPCs through three successive rounds of incubation on ice and centrifugation at 10,000 x g at 4 °C. The supernatants were removed each time. Upon completion of the last washing step, the final pellets were re-suspended in 1 mL DPC pellet digestion Buffer [100 mM KCl, 10 mM EDTA, 20 mM Tris-HCl (pH 7.5) with 300 μg/mL proteinase K]. 50 μL (equal volume) of Input digestion buffer [200 mM KCl, 10 mM EDTA, 20 mM Tris-HCl with 30 μg/mL proteinase K] was added to the “Input DNA fractions” samples. After overnight incubation of both “DPC pellets” and “Input DNA” fractions at 37 °C, samples were centrifuged at 20,000 × g for 10 min. The supernatants from both fractions were collected and transferred to new Eppendorf tubes for DNA quantification. The total DNA amounts were determined using the Promega QuantiFluor dsDNA system kit according to the manufacturer’s instructions. DPC levels were calculated as the ratio of DNA amount from the “DPC pellet fractions” to the DNA from the “Input DNA fractions” and reported as a percentage of DNA containing DPCs.

### DNA quantification using Promega QuantiFluor dsDNA Assay

DNA amounts in the input and DPC fractions yields were determined using a Promega QuantiFluor dsDNA assay kit according to the manufacturer’s procedure. In brief, 200 μL of prepared QuantiFluor ONE dsDNA dye solution was added to wells designated for unknown samples and standards in 96-well plates. Next, dsDNA standards for the standard curve were prepared by serially diluting the QuantiFluor ONE lambda DNA stock (400 ng/μL). After adding equal volumes of both standards and unknown samples to the dye solution, the plate was incubated in the dark for 5 minutes, and fluorescence was measured at 504 and 531 nm wavelengths using a BioTek Synergy H1 plate reader.

### Quantification of DPCs in H_2_O_2_-treated cells

Human fibrosarcoma (HT1080) cells (1.2 × 10^6^ cells/mL), cultured in DMEM media for 72 h, were treated with 200 μM H_2_O_2_ for 1 h. To characterize ROS-DPC formation in mammalian cells exposed to H_2_O_2_, DPC-containing DNA was isolated using the published phenol-chloroform extraction method (5) with a few modifications. Briefly, after a 1-hour 200 μM H_2_O_2_ treatment, HT1080 cells were lysed in a 2X lysis buffer [20 mM Tris-HCl (pH 7.4), 10 mM MgCl_2_, 2% Triton X-100, and 0.65M sucrose], and solubilized with 22G syringes. Following a 10 min centrifugation at 2,000 x g at 4 °C, nuclear pellets were dissolved in 2 mL of a saline-EDTA solution [75 mM NaCl, 24 mM EDTA, 2% SDS (pH 8.0), 10 μg/mL RNase A, and complete protease inhibitor cocktail tablets (EDTA-free) (Thermo fisher scientific; 1 tablet per 10 mL buffer) for 2 hours at 37 °C. After three successive additions of a 1:1 mixture of phenol and chloroform, protein-bound DNA was isolated from both interface and aqueous layers. DPC-containing DNA was ethanol-precipitated overnight at −20 °C, washed, and reconstituted in 300 μL re-suspension buffer [20 mM Tris-HCl (pH 7.4), 24 mM EDTA]. A small portion of extracted DPCs (50 μL) was digested with 30 µg/mL proteinase K, and DNA was quantified using dG quantitation by HPLC-UV as described below. Protein levels were quantified using the Pierce Bradford Assay per the manufacturer’s protocol.

### DNA quantitation by HPLC-UV analysis of 2-deoxyguanosine (dG)

DPC-containing DNA was quantified using an HPLC-UV-based dG assay as described previously (24). In brief, 2 µg aliquots of DNA (Nanodrop) were digested to nucleosides with the following enzyme mix: phosphodiesterase I (120 mU), phosphodiesterase II (105 mU), DNase I (35 U), and alkaline phosphatase (22 U) in digestion buffer [10 mM Tris-HCl (pH 7.0), and 15 mM ZnCl_2_]. The reaction was left for 18 hours at 37 °C. Enzymes were removed using *Nanosep* 10K filters and centrifuged at 10,000 × g for 5 min. dG was quantified using HPLC-UV on an Agilent Technologies 1100 HPLC system. HPLC was equipped with an autosampler and a diode array UV detector. Following their transfer to HPLC vials, the samples were loaded onto an Atlantis T3 C18 column (0.21 x 15 cm, 5 *μ*m, Waters Corporation, Milford, MA). A gradient of solvent A [0.005M ammonium formate (pH 4.0)] and solvent B [100% methanol] was used to separate DNA nucleosides. The fraction of the organic phase was increased linearly from 3% to 30% B over 15 min, then raised to 80% B over 3 min, maintained at 80% B for 1 min, and lowered to 3% B over 2 min, where it remained for the last 8 min of the HPLC run. With this method, dG eluted as a sharp peak at ~11.59 min **(Figure S1)**. dG amounts were calculated by comparing HPLC-UV peak areas corresponding to the dG to the standard curve generated by injecting known dG amounts.

### Rapid Approach to DNA Adduct Recovery Assay (RADAR)

DNA-crosslinked proteins were isolated using a modified Rapid Approach DNA Adduct Recovery (RADAR) assay (56). In brief, HT1080 cells (2 million), treated with 0, 0.2, 0.5, 1.0 mM H_2_O_2_ for 2 h, were lysed in 1 mL M buffer [6M guanidine isothiocyanate, 10 mM Tris-HCl (pH 6.8), 20 mM EDTA, 4% Triton X-100, 1% Sarkosyl, and fresh 1% dithiothreitol]. DNA was precipitated by adding 1 mL of 100% ethanol to the samples and washed three times with aqueous solution containing 20 mM Tris-HCl, pH 6.8, 150 mM NaCl, and 50% ethanol. DNA was sheared using 22G syringe needles and sonicated (60% amplitude, 15 seconds, 1s on, 1s off pulses) to facilitate solubilization in 400 μL TE buffer [10 mM Tris-HCl (pH 6.8), and 1 mM EDTA]. After reversing crosslinks at 70 °C for 1 h, samples were treated with RNase A (10 μg/mL) for 15 min at 37 °C. A small aliquot of the resulting DNA was digested with 50 µg/mL proteinase K (Invitrogen) at 50 °C for 3 h and quantified using the Promega Quant-Fluor dsDNA System Assay Kit. Equal amounts of DNA were loaded onto 4-12% Bis-Tris SDS-PAGE gels, followed by Western blotting for ubiquitin. Equal amounts (~30 ug) of DNA were digested with 5U nuclease P1 enzyme and loaded onto 4-12% Bis-Tris SDS-PAGE gels, followed by western blotting for ubiquitin.

### SDS-PAGE separation and staining of DNA-crosslinked proteins

To separate and visualize DNA-crosslinked proteins, 10 μg aliquots of DPCs extracted via the modified phenol-chloroform method described above were sonicated using Fisherbrand™ model 120 sonic dismembrator (with CL-18 probe) (60% amplitude, 15-second pulse, 1 sec on, 1 sec off). Samples were then digested with 5 U nuclease P1 overnight at 37 °C to remove DNA. The next day, the samples were heated at 90 °C in denaturing NuPAGE LDS buffer (Thermo Scientific; Buchs, Switzerland) for 10 min, vortexed for 5 min, and loaded onto 4-12% NuPAGE SDS-PAGE gels (Thermo Fisher Scientific; Waltham, MA). DPC proteins were separated for 90 min at 100 V. Gels were washed in Milli-Q water, stained with SimplyBlue stain for 2 h and destained overnight in Milli-Q water.

### Western blot analyses

To confirm the identities of ROS-DPC participating proteins, DPC containing DNA (30 μg) was digested with nuclease P1 for 4 h at 37 °C. The liberated proteins were separated using 4-12% Bis-Tris SDS-PAGE gels. Following separation, proteins were transferred to nitrocellulose membranes (Bio-Rad, Hercules, CA). Membranes were blocked for 1 h at room temperature in Tris-buffered saline (TBS) containing 2.5% dry milk and 0.1% Tween-20. Membranes were incubated overnight at 4 °C with primary antibodies against ubiquitin (Catalog No. 43124T, Cell Signaling Technology; 1:10000). After three 5 min washes with TBS supplemented with 0.1% Tween-20 detergent (TBST, ICN Biomedicals), membranes were incubated with secondary antibodies (LiCor, at 1:10,000 dilution) for 1 hour at RT. Following three washes, membranes were scanned using a LI-COR Odyssey FC instrument.

### Mass spectrometric identification of DNA-crosslinked proteins

#### Sample preparation for label free proteomics

To identify cellular proteins that participate in DPC formation in H_2_O_2_-treated cells, 30 µg of DPC-containing DNA isolated by the phenol-chloroform method was dissolved in 100 μL of 50 mM ammonium acetate buffer (pH 5.5) supplemented with 10 μL of 5 mM ZnCl_2_. Samples were then heated at 90 °C for 10 min and plunged in ice for 20 min to denature DNA. Two units of nuclease P1 were added to the chilled samples, and the DNA was digested for 4 h at 37 °C. The pH of each sample was adjusted to 7.0 by adding 1 μL of 1 M ammonium bicarbonate, and the samples were further digested overnight by adding 120 U phosphodiesterase I and 22 U alkaline phosphatase (Sigma, St. Louis, MO). The next day, samples were transferred to Amicon 3K filters (Millipore, Darmstadt, Germany), and free nucleosides were removed by centrifugation at 14,000 × g for 10 minutes. The filtrates were discarded, and the protein-nucleoside conjugates were washed on filter three times with 100 mM HEPES (pH 8) and re-dissolved in a reduction buffer [0.1M NaPO_4_, 1mM EDTA (pH 8)]. A 10 μL aliquot of 100 mM DTT/0.1% SDS/100 mM HEPES (pH 8) was added to each sample, and the samples were heated at 55 °C for 1 h in the dark. Next, 5 μL of 375 mM iodoacetamide was added to alkylate reduced cysteine residues, and the samples were incubated for 30 min at 25 °C. Excess reagents were removed by centrifuging the samples at 14,000 g for 10 min. Proteins were resuspended in 50 μL of 25 mM ammonium bicarbonate, and 8 U of Pierce™ Lys-C/trypsin enzyme mixture (Thermo Scientific) was added to digest proteins to peptides overnight at 37 °C. The resulting tryptic peptides were resuspended in a 95:5:0.1 acetonitrile: water: formic acid solution prior to nanoHPLC-ESI^+^-MS/MS analysis.

#### NanoHPLC-ESI^+^ −MS/MS analyses of tryptic peptides

To identify and quantify DPC proteins, peptide samples were analyzed on an Eksigent NanoLC-Ultra 2D HPLC interfaced to a nanospray ESI source and Q Exactive MS (Thermo Scientific, Waltham, MA). The injection volume was 5 μL. Samples were separated on an in-house-packed C-18 column (15 cm × 75 μm, Luna C-18 resin, Phenomenex, Torrance, CA) using a 60-min method gradient. HPLC flow rate was maintained at 500 nL/min. 0.1% formic acid in water was the mobile phase solvent A, while 0.1% formic acid in acetonitrile was the solvent B. Solvent composition was held at 5% solvent B for 5 min, followed by an increase to 35% over 40 min, further to 95% over 9 min, returned to 5% over 3 min, and re-equilibrated for 30 min. Typical ion source parameters were set to a spray voltage of 3000 V, a capillary temperature of 300 °C, and a 50% S-lens RF level. MS analysis was performed in the data-dependent mode with dynamic exclusion enabled (1 repeat count and exclusion time of 10 seconds). The 12 most intense precursor ions were selected for HCD fragmentation, which was set to 30%. Every full MS scan (*m/z* 350-1800) was collected at a resolution of 70,000, with an AGC target of 1 × 10^6^ and an isolation window of 1.8 m/z.

#### Bioinformatic analysis of mass spectrometry results

Raw MS files were searched against the human proteome (UP000005640, retrieved on 2024-10-06 from UniProt), decoys, and common contaminants with Flagpipe (version 22.0, MSFragger version 4.1, IonQuant version 1.10.27, Python version 3.9.13), using the default settings of the label-free quantification-match between runs (LFQ-MBR) workflow. Within the software, peptides were matched to the spectra and assigned to proteins with an FDR of 1%. Missing values in the tabular output file of protein data were imputed so that zeros could be replaced with half the minimum non-zero MaxLFQ intensity value. The log2-transformed *p*-values were calculated using Prism (version 9.4.1) using unpaired, parametric t-tests.

## RESULTS

### DPC formation in vitro and in H_2_O_2_-treated human cells

To determine whether H_2_O_2_ can induce DPC formation *in vitro*, we employed a model system consisting of a 69-mer biotinylated double-stranded oligonucleotide and recombinant HMGB2, a high-mobility group DNA-binding protein located in the nuclei of eukaryotic cells that is involved in transcription, chromatin remodeling, and immune responses (57). HMGB2 protein (M.W. 28 kDa) was incubated with synthetic DNA duplex (Biotin-5’-TTG GTA AAA CGA CGG CCA GTG CCA AGC CAT GCC TGC AGG TCG ACT CTA GAG GAT CCC CCA GCC TGC CGG-3’, M.W. 43 kDa) in the presence of 0, 0.1, 1, and 5 mM H_2_O_2_. Denaturing PAGE analysis of the reaction mixtures revealed a concentration-dependent formation of a new high-molecular-weight species with a molecular weight of 71 kDa corresponding to a covalent conjugate between HMGB2 and the DNA duplex **(Figure 1A)**. Additional high-molecular-weight bands at 125 kDa and above were observed, perhaps due to additional cross-linking events.

**Figure 1.**
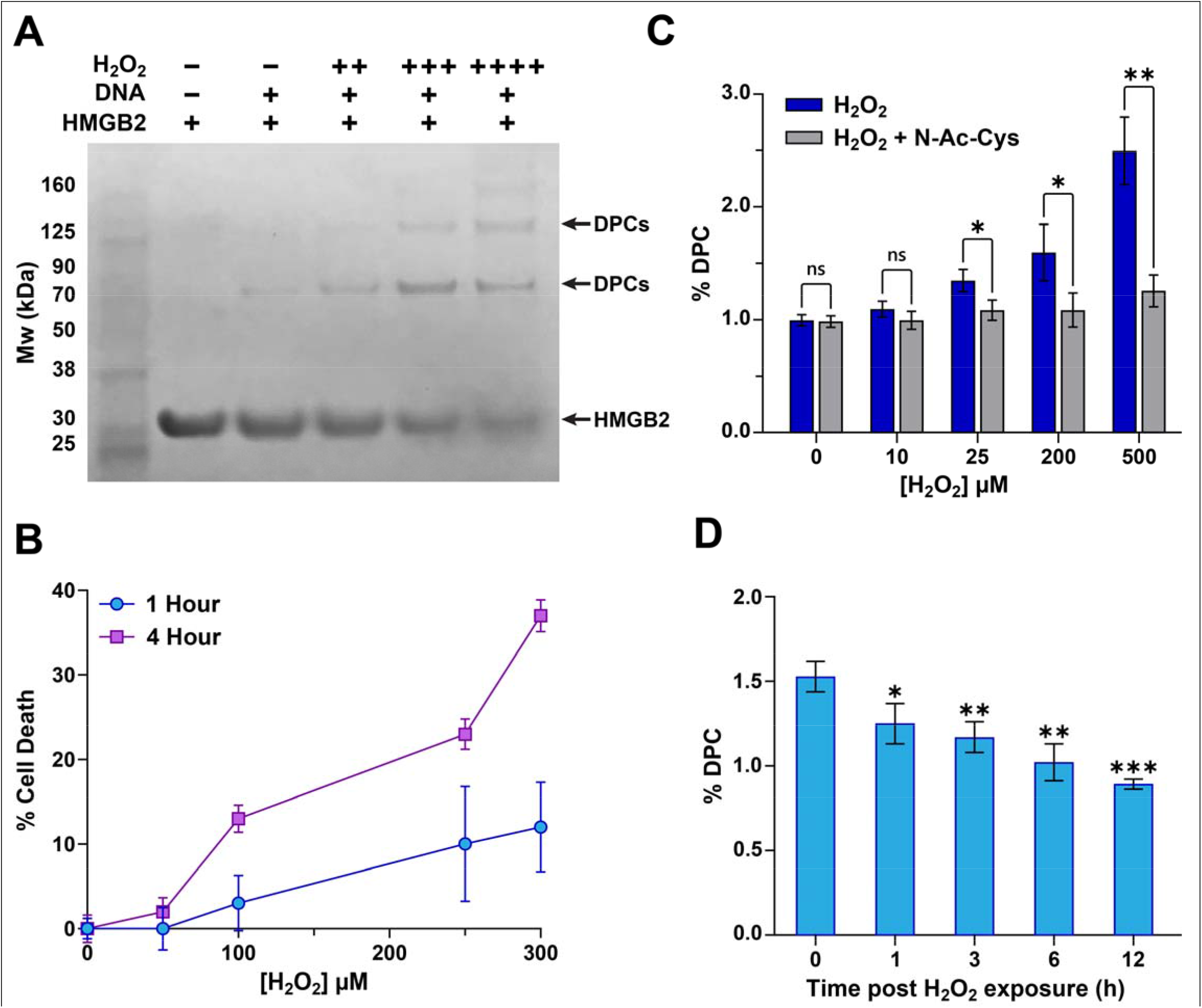
H_2_O_2_ exposure induces DNA-protein crosslinks (DPCs). **(A)** *H*_2_*O*_2_ *induces DPCs in vitro*. Recombinant high mobility group box 2 protein (HMGB2) (500 pmol) was incubated with biotinylated double-stranded a 69-mer DNA duplex (250 pmol) in the absence or in the presence of H_2_O_2_ (0.1,1 or 5 mM) in 100 mM NH_4_CO_3_ (pH 8) for 3 h at 37 °C. Proteins and DPCs were separated using the 4-12% NuPAGE SDS-PAGE gel and stained with SimplyBlue stain. **(B)** *H*_2_*O*_2_ *induces cell death*. HT1080 cells were treated with 50, 100, 250, or 300 *μ*M H_2_O_2_ for 1 or 4 h. Cell death as a function of time and concentration was determined using the alamar blue cell viability assay. **(C)** *N-acetyl-L-cysteine ((NAC) blocks H*_2_*O*_2-_*induced DPCs in cells*: HT1080 cells were pre-treated with N-Ac-Cys) a day prior to H_2_O_2_ exposure at indicated concentrations, followed by the K-SDS assay to quantify DPC levels. DPCs were quantified using the Promega QuantiFluor dsDNA assay. Data were analyzed via a one-way ANOVA and an unpaired t-test (\**p* < 0.05, *: *p*-value < 0.05, **: *p*-value < 0.01, ***: *p*-value < 0.001). Values are reported as mean ± standard deviation (SD) from three independent biological replicates. **(D)** *DPC repair time course in HT1080 cells*. Cells were exposed to 200 *μ*M H_2_O_2_ for 1 h, followed by quenching with excess N-Ac-Cys. Cells were washed with PBS and placed in fresh media for the indicated times. DPCs were quantified via the K-SDS assay. DPC levels at the end of designated recovery times were compared to the DPC level at T0 (DPCs in treated cells). Data were analyzed via a one-way ANOVA and an unpaired t-test (\**p* < 0.05, *: *p*-value < 0.05, **: *p*-value < 0.01, ***: *p*-value < 0.001). Values are reported as mean ± standard deviation (SD) from three independent biological replicates.

To evaluate H_2_O_2_ cytotoxicity, HT1080 cells were exposed to increasing concentrations (0-300 μM) of H_2_O_2_ for 1 or 4 h, followed by a 17-hour recovery period. Cell viability was assessed using the alamar blue assay. H_2_O_2_ treatment led to time and concentration-dependent increase in cell death, with 4 h exposure resulting in higher toxicity overall (**Figure 1A**). Treatment with 300 μM H_2_O_2_ for 1 or 4 h induced cell death amounting to 12 ± 5.0% and 37 ± 2.0%, respectively (**Figure 1A)**.

To examine DPC formation in cells upon H_2_O_2_ treatment, we employed the K-SDS assay (7,58). In this approach, cells are lysed and denatured with sodium dodecyl sulfate (SDS). Potassium chloride (KCl) is added to cell lysates, precipitating any protein-bound DNA while leaving free DNA in solution. Any RNA-protein crosslinks are removed via digestion with RNase A. As shown in Figure 1C, DPC levels in HT1080 cells rose from 1.00% ± 0.05% in untreated cultures to 1.60% ± 0.25% in cells exposed to 200 μM H_2_O_2_, and further to 2.50% ± 0.30% in cells treated with 500 μM H_2_O_2_, confirming the formation of ROS-induced DPC in cells treated with hydrogen peroxide. Consistent with our previous study (5), significant levels of endogenous DPCs (1 %) were observed in untreated cells. These background DPCs may arise from exposure to electrophilic cellular metabolites such as formaldehyde and methylglyoxal (18,59), endogenous ROS, or could be formed enzymatically when topoisomerases, DNA repair proteins, and DNA polymerases are trapped on DNA during catalysis (16,17). Future studies are needed to characterize endogenously formed DPCs.

N-acetyl-L-cysteine (N-Ac-Cys) is an antioxidant and a reducing agent that can inactivate cellular ROS (60). When HT1080 cells were pre-treated with N-Ac-Cys and exposed to H_2_O_2_ on the following day, DPC levels in cells treated with 500 μM H_2_O_2_ dropped from 5.0% ± 0.80% to 1.3% ± 0.20% **(Figure 1C)**. These results indicate that DPC formation in human cells can be reduced by antioxidant treatment, suggesting a possible therapeutic strategy.

To analyze the persistence of ROS-induced DPCs in human cells, HT1080 cells were treated with 200 μM H_2_O_2_ for 1 h. H_2_O_2_ was quenched with N-Ac-Cys, followed by re-plating and recovery in fresh media for 1-6 h. DPC levels post exposure were monitored by the K-SDS assay described above. Cellular ROS-DPC levels returned to the endogenous levels (1.0% ± 0.11%) after 6 h of recovery **(Figure 1D)**, with the calculated half-life (t_1/2_) of 2 h. These results indicate that H_2_O_2_-induced DPCs are actively repaired in human cells.

### Identification of DNA-crosslinked Proteins by Mass Spectrometry-Based Proteomics

To isolate the proteins involved in H_2_O_2_-induced DPC formation, we employed a modified phenol-chloroform extraction methodology (5,59,61). This assay relies on phase partitioning, in which proteins covalently bound to genomic DNA migrate to the interface between the aqueous and the organic phase, while free proteins are extracted with organic solvent (5,59,61). Following extraction, DPCs isolated from treated cells were digested with nuclease P1 to remove DNA (**Figure 2A**). The liberated proteins were separated on a 4-12% SDS-PAGE gel, followed by visualization with Simply Blue staining. Distinct protein bands spanning the entire range of molecular weights were observed for H_2_O_2_-treated samples, whereas untreated samples showed only weak protein band signals (48.8 AU versus 3.1 AU, **Figure 2B**). The remaining samples were subjected to tryptic digestion and label-free label free nanoLC-NSI-MS/MS analyses to identify proteins involved in DPC formation.

**Figure 2.**
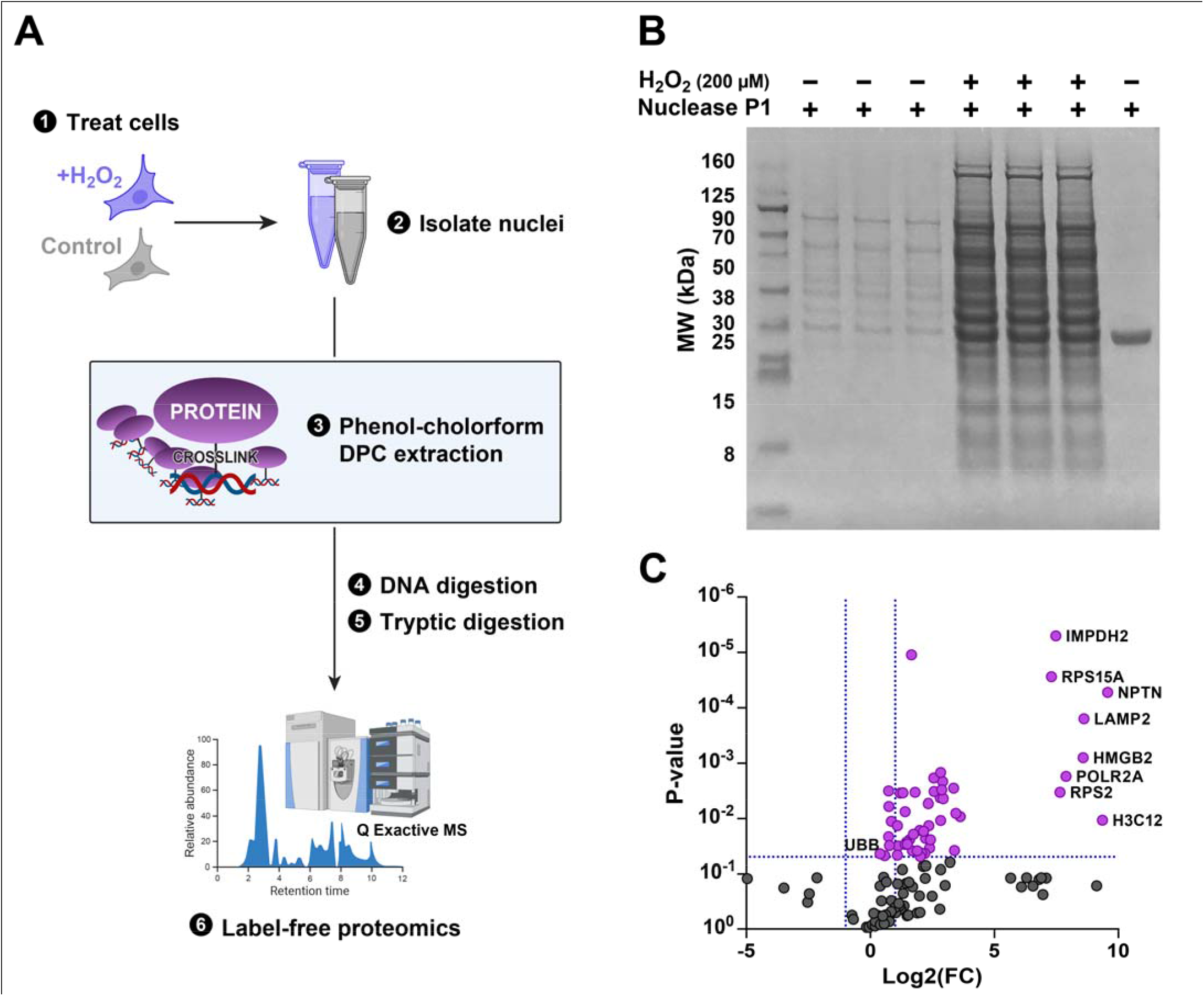
SDS-PAGE analysis and characterization of H_2_O_2_-derived DPCs in HT1080 cells by label free mass spectrometry-based proteomics. **(A)** *Experimental scheme for identifying DNA-crosslinked proteins*. **(B)** *SDS-PAGE separation of proteins involved in DPC formation by H*_2_*O*_2_. DPC-associated proteins were isolated from H_2_O_2_-treated using modified phenol-chloroform assay as described in the materials and methods section. DNA was removed via digestion with Nuclease P1 prior to loading samples on 4-12% NuPAGE SDS-PAGE gel and staining with SimplyBlue stain. **(C)** *Label free proteomics results for H*_2_*O*_2_*-induced DPCs*. A volcano plot illustrating proteins that participate in H_2_O_2_-induced DPC formation (*n=3* independent experiments for H_2_O_2_-treated samples and their vehicle controls) as identified by label-free MS-based proteomics. Proteins represented by magenta-colored dots were significantly enriched in H_2_O_2_-treated samples (*p*-values <0.05).

Spectral data from label free nanoLC-NSI-MS/MS analyses of DPC peptides were analyzed using the Flagpipe computational platform, where peptides were searched against a human proteome FASTA file retrieved from the UniProt database (www.uniprot.org). These analyses have identified 105 proteins that showed a statistically significant increase in DPCs in HT1080 cells treated with H_2_O_2_. These proteins were enriched in the H_2_O_2_ treatment group (*p*-value < 0.05).

**Table S1** lists all of the identified DPC-proteins, including known DNA binders such as high-mobility group proteins (HMG), RNA polymerase, and histone proteins, which have been previously reported in other DPC studies (**Figure 2C**) (5,24,61). The DAVID software was employed to elucidate the biological processes and cellular compartments of identified proteins (**Figure S2**). The majority of the proteins involved in ROS-mediated DPC formation have roles in structural and functional processes involving DNA including nucleosome assembly, gene expression, and telomere organization. These analyses also identified several proteins involved in RNA-dependent processes. Since our workflow employed an RNase digestion step, any RNA-protein conjugates should have been removed. This was confirmed by HPLC-UV analyses of hydrolysates, demonstrating the absence of RNA nucleosides **(Figure S1A)**. Therefore, we hypothesize that these known RNA binders also exhibit DNA-binding activity, allowing them to be crosslinked to DNA. This is consistent with our previous proteomics studies, which also found RNA-binding proteins enriched among DNA-crosslinked targets (5,62).

### SPRTN and the Ubiquitin-proteasomal system (UPS) constitute major proteolytic pathways involved in the removal of ROS-derived DPCs

Initial steps of DPC repair require proteolytic processing of the protein component of DPCs. Both SPRTN metalloprotease and ubiquitin-proteasomal system (UPS) have been previously implicated in DPC proteolysis (28,36,37). To examine whether ROS-DPCs are processed by SPRTN, we employed Sprtn^f/-^ mouse embryonic fibroblasts (MEF5) with an intact floxed allele, which exhibit significantly reduced SPRTN gene expression (54). Diminished expression of SPRTN resulted in increased cell death following H_2_O_2_ treatment (**Figure 3A**). Additionally, Sprtn^f/-^ MEFs accumulated significantly higher levels of H _2_O_2_-derived DPCs than their wildtype counterparts **(Figure 3B)**. Finally, SPRTN deficiency also increased the levels of endogenous DPCs in cells that were not subjected to any treatment **(Figure 3B)**. This is consistent with our previous observation of elevated levels of Thy-Tyr DPCs in tissues of unexposed SPRTN-deficient mice (7). Collectively, these finding provide additional evidence that SPRTN metalloprotease is involved in the removal of ROS-induced DPCs in human cells.

**Figure 3.**
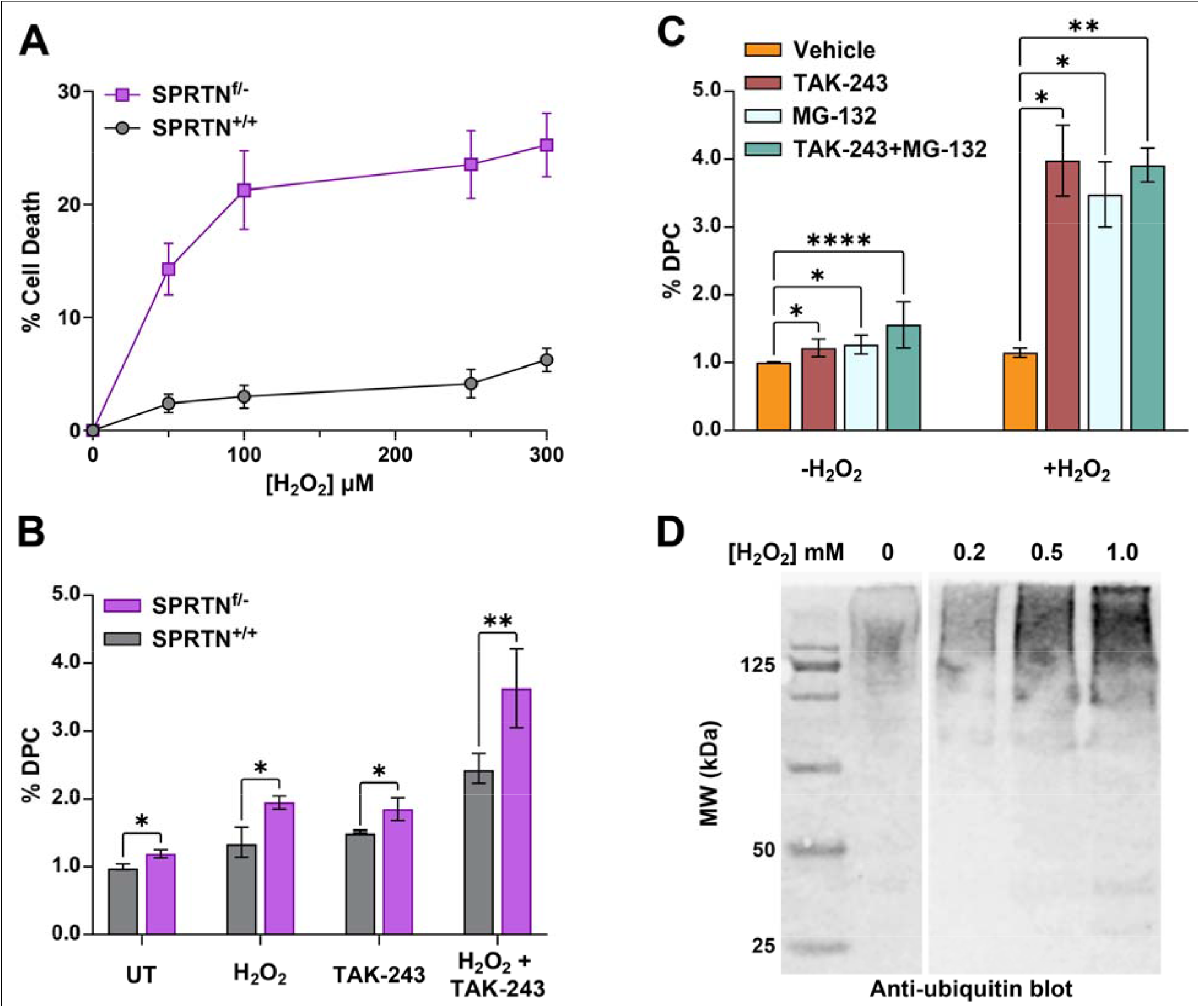
SPRTN and the Ubiquitin-proteasomal system (UPS) constitute major proteolytic pathways involved in the removal of ROS-derived DPCs. **(A)** *SPRTN deficient cells are sensitized to ROS*. MEFs were exposed to H_2_O_2_-containing media (0, 50, 100, 250, and 300 µM H_2_O_2_). Viability was determined using an alamar blue assay. % Cell deaths were computed by comparing cell count in treatment conditions relative to the untreated controls. Data is represented as mean ± SD from three biological replicates. **(B)**. *SPRTN is involved in ROS-DPC repair and requires DPC ubiquitination*. Untreated and H_2_O_2_-treated MEFs (0 or 200 µM for 1h) before or after pre-treatment with the inhibitor of ubiquitin activating enzyme 1, TAK-243 (10 µM), were processed using a K-SDS assay to isolate DPC-associated DNA. Data are represented as mean ± SD from three biological replicates. Analysis was performed via a one-way ANOVA and an unpaired t-test (\**p*-value < 0.05). The asterisks indicate t-test *p*-values from one-way ANOVA with \**p*-value < 0.05, \*\**p*-value < 0.01, \*\*\**p*-value < 0.001, and \*\*\*\**p*-value < 0.00001. **(C)** *Ubiquitin-Proteasomal pathway facilitates ROS-DPC removal in human cells*. HT0180 cells were pre-treated with 10 µM TAK243, MG132 (26S proteasomal subunit inhibitor), or their combination in the presence of 0 or 200 µM H_2_O_2_ for 1 h before H_2_O_2_ treatments. Following the H_2_O_2_ quenching and a second PBS wash of the cells, DPC-associated DNA was isolated immediately using a K-SDS assay and quantified using Promega QuantiFluor dsDNA assay to assess DPC levels. Data are represented as mean ± SD from three biological replicates. Analysis was performed via a one-way ANOVA and an unpaired t-test (\**p*-value < 0.05). The asterisks indicate t-test *p*-values from one-way ANOVA with \**p*-value < 0.05, \*\**p*-value < 0.01, \*\*\**p*-value < 0.001, and \*\*\*\**p*-value < 0.00001. **(D)** *Western blot analysis confirms ubiquitination of H*_2_*O*_2_*-induced DPCs in HT1080 cells*. Following treatment of the HT1080 cells with 0, 0.2, 0.5, and 1 mM H_2_O_2_ for 1 h, and DPCs were extracted by the RADAR assay. Samples were normalized by DNA amount. Proteins from 30 µg of DNA were digested with Nuclease P1, separated by SDS-PAGE gels and transferred to nitrocellulose membranes. Western blotting was performed using a primary antibody against ubiquitin.

DPC processing may be facilitated by ubiquitination of the DPC proteins. Ubiquitin serves as a recognition signal to recruit SPRTN metalloprotease to DPC site via its ubiquitin-binding domain (28,36,37). Additionally, polyubiquitination of DPC proteins allows for their degradation by the proteasome (63–65) Therefore, we investigated whether the E1 ubiquitin ligase pathway facilitates the removal of H_2_O_2_-derived DPCs. Pre-treatment of Sprtn^f/-^ MEFs with TAK-243, an inhibitor of ubiquitin activating enzyme 1 (UAE1), led to a dramatic increase of DPC numbers in H_2_O_2_-treated cells (3.6% ± 0.58%) and a smaller effect in untreated control (**Figure 3C**). Western blotting confirmed ubiquitination of DNA-crosslinked proteins **(Figure 3D)**.

Polyubiquitinated proteins, including DPC proteins, can be cleaved to peptides by the ubiquitin-proteasomal system (UPS) (50,66). To determine whether the UPS plays a role in the cellular repair of H_2_O_2_-derived DPCs, we co-treated HT1080 cells with 200 *μ*M H_2_O_2_ and pharmacological inhibitors of the ubiquitin-activating enzyme (UAE1), TAK-243, and/or inhibitor of the 26S proteasomal subunit (MG-132). Each inhibitor was sufficient to cause a significant accumulation of both endogenous and H_2_O_2_-derived DPCs, and co-treatment with both TAK243 and MG132 led to even higher DPC levels in human cells (Figure 3C). Exposure of HT1080 cells to both TAK-243 and MG-132 led to a 3-fold increase in H_2_O_2_-derived DPCs (3.9% ± 0.25%) compared to treatment with TAK-243 (1.2% ± 0.13%) or MG-132 (1.3% ± 0.14%) only **(Figure 3C)**. These data indicate that proteolytic processing of H_2_O_2_-derived DPCs requires ubiquitination activity and that, in addition to SPRTN metalloprotease, the proteasome plays an essential role in ROS-DPC removal.

### Cellular replication, transcription, and NER pathways play a role in clearance of H_2_O_2_-DPCs

Since DNA-protein interactions guide most of the biological processes involving DNA, cellular repair machinery must distinguish between covalent and noncovalent DNA-protein complexes (DPCs). Human cells could employ the transcription and replication machineries as surveillance mechanisms to detect DPC lesions and to initiate DNA damage response (27,67). To test whether transcription and replication play a role in detection and/or removal of H_2_O_2_-DPCs, we co-treated H1080 cells with H_2_O_2_ and pharmacological inhibitor of replication, aphidicolin (APH), or transcription inhibitor, actinomycin D (Act D), followed by K-SDS measurements of DPC amounts. Treatment with APH and Act D induced about a 2-fold increase in endogenous DPCs, and a 3-fold increase in the H_2_O_2_-derived DPCs. Act D/H_2_O_2_; 5.24 ± 0.06% DPCs vs APH/H_2_O_2_; 3.59 ± 0.15% DPCs) **(Figure 4A)**. These results indicate that mammalian cells repair ROS-DPC lesions via replication- and/or transcription-dependent manner.

**Figure 4.**
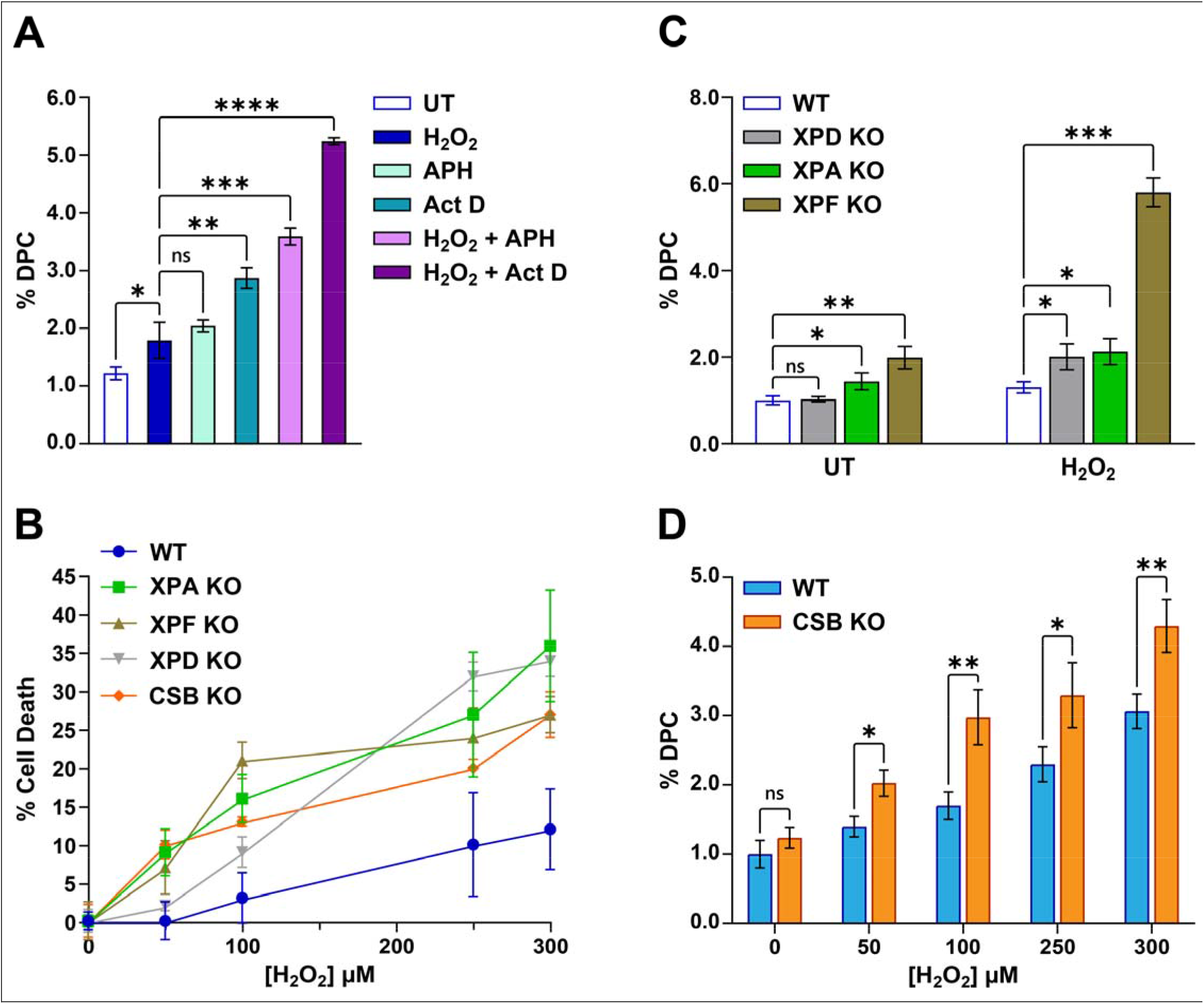
Role of cellular replication, global-genome NER and transcription-coupled NER machineries in the clearance of ROS-DPCs. (A). *Replication and transcription facilitate the repair of ROS-DPCs*. HT1080 cells were pre-treated with 10 *μ*M of the replication inhibitor APH or the transcription inhibitor Act D for 24 h prior to H_2_O_2_ exposure (200 *μ*M for 1 hour). DPCs were detected by the K-SDS assay. Data are represented as mean ± SD from three biological replicates. Analysis was performed via a one-way ANOVA and an unpaired t-test (\**p*-value < 0.05). The asterisks indicate t-test *p*-values from one-way ANOVA with \**p*-value < 0.05, \*\**p*-value < 0.01, \*\*\**p*-value < 0.001, and \*\*\*\**p*-value < 0.00001. **(B)** *Deficiency in NER factors sensitizes cells to ROS-induced death*. XPA-, XPF-, XPD-, and CSB-deficient and wild-type HT1080 cells were treated with H_2_O_2_ at the indicated concentrations for 1 h, followed by 17 h recovery. Cell viability was assessed using the Alamar blue assay. Data are presented as mean ± SD from 3 replicates and normalized to a vehicle control. **(C)** *NER deficiency leads to DPC accumulation in H*_2_*O*_2_ *treated cells*. ROS-DPC formation was examined by treating XPA, XPF-, XPD-, CSB-deficient, and WT HT1080 cells with 200 *μ*M H_2_O_2_ for 1 h. DPCs were quantified using the K-SDS assay. Data are represented as mean ± SD from three biological replicates. Analysis was performed via a one-way ANOVA and an unpaired t-test (\**p*-value < 0.05). The asterisks indicate t-test *p*-values from one-way ANOVA with \**p*-value < 0.05, \*\**p*-value < 0.01, \*\*\**p*-value < 0.001, and \*\*\*\**p*-value < 0.00001. **(D)** *TC NER is involved in ROS-DPC removal*. CSB KO and wild-type HT1080 cells were treated with 0, 50, 100, 250, and 300 *μ*M H_2_O_2_ for 1 h. DPCs were quantified using the K-SDS assay and quantified using the Promega QuantiFluor dsDNA assay. Data are represented as mean ± SD from three biological replicates. Analysis was performed via a one-way ANOVA and an unpaired t-test (\**p*-value < 0.05). The asterisks indicate t-test *p*-values from one-way ANOVA with \**p*-value < 0.05, \*\**p*-value < 0.01, \*\*\**p*-value < 0.001, and \*\*\*\**p*-value < 0.00001.

Following proteolytic digestion of DNA-bound proteins, canonical DNA repair pathways such as nucleotide excision repair (NER), can be used to repair the resulting DNA-peptide cross-links (DpCs). Transcription-coupled NER (TC-NER) is active in transcribed regions of the genome, while global genome NER (GG-NER) surveils the entire genome for bulky/distorting DNA lesions (68,69). To investigate the role of the NER pathways in removal of H_2_O_2_-derived DNA lesions, including DPCs, we employed repair deficient clones HT1080-XPA, XPD, XPF, and CSB. XPA, XPD, and XPF are all central to both gg-NER and TC-NER, while CSB binds to RNA polymerase II as an exclusive TC-NER recognition factor. Total loss of these NER factors sensitized HT1080 cells to H_2_O_2_ cytotoxicity **(Figure 4A)** and led to significantly increased DPC levels **(Figure 4B&C)**, confirming the involvement of NER in H_2_O_2_-DPC removal.

To further assess the role of gg-NER and TC-NER factors in the clearance of H_2_O_2_-DPCs, we compared DPC levels in wildtype HT1080 cells and NER-deficient clones treated with H_2_O_2_. XPA- and XPD-deficient cells (green and gray) showed approximately 2-fold increase in H_2_O_2_-DPCs, while the loss of XPF (brown) resulted in the highest levels of DPCs (6%) **(Figure 4C)**. Similarly, CSB KO cells exhibited significantly higher levels of H_2_O_2_-DPCs than wild-type (WT) cells, with a maximum of 4.30 ± 0.38% DPCs at 300 μM H_2_O_2_ treatment (vs 3.07% ± 0.25% DPCs from WT cells) **(Figure 4D)**. In contrast, inhibiting the homologous repair pathway with RAD51 recombinase inhibitor B02 had no effect on H_2_O_2_-DPC levels **(Figure S3)**. Both NER and HR pathways were previously shown to participate in the removal of other types of DPCs, such as formaldehyde and decitabine-induced DPCs (42,70). Similarly, ROS-DPC levels were unaffected by inhibiting autophagy with chloroquine **(Figure S3)**.

### Synthetic glutathione analog (Ψ-GSH) attenuates H_2_O_2_-DPC formation in human cells

Glutathione (GSH) plays a key role in scavenging physiological ROS and maintaining a reducing environment in cells (71). However, native GSH is rapidly cleaved by y-glutamyl transpeptidase. We have recently reported that GSH analogue (Ψ-GSH), which contains a ureide linkage instead of γ-glutamylcysteine amine linkage **(Figure 5A)**, is stable *in vivo*, enters the brain, and attenuates the symptoms of Alzheimer’s disease in a mouse model (72,73). To determine whether Ψ-GSH can prevent H_2_O_2_-DPC formation, HT0180 cells were pre-treated with increasing concentrations of Ψ-GSH prior to exposure to 200 *μ*M H_2_O_2_ for 1 h. Ψ-GSH treatment resulted in a concentration-dependent reduction of levels of H_2_O_2_-DPCs and also reduced the levels of endogenous DPCs in untreated cells (**Figure 5B**). These results are consistent with our results for N-acetylcysteine **(Figure 1C)** and demonstrate the ability of the Ψ-GSH to prevent the formation of ROS-DPCs in living cells.

**Figure 5.**
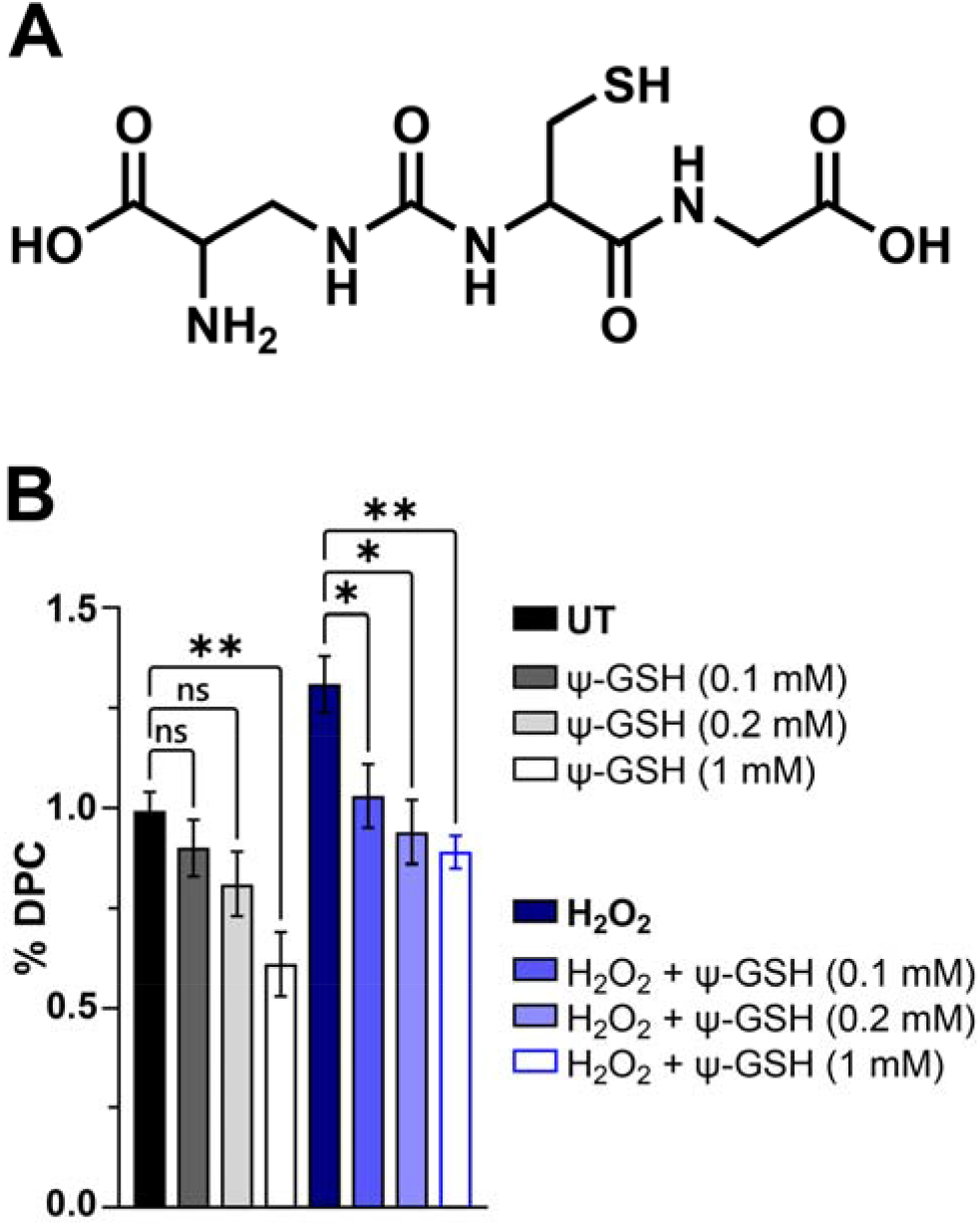
The γ-Glutamyl-transpeptidase-resistant glutathione analog, Ψ-GSH, attenuates H_2_O_2_-induced DPC formation. **(A)** Chemical structure of Ψ-GSH (72). **(B)** HT1080 cells were pre-treated with increasing GSH concentrations (0.1, 0.2, and 1 mM) for 24 h before a 1 h treatment with 200 **μ**M H_2_O_2_. DPCs were quantified via the K-SDS assay. Data are presented as the mean ± SD values of three independent experiments. Ψ-GSH treatment groups were compared with either the untreated control or the H_2_O_2_ groups using one-way ANOVA and unpaired t-test analyses. Asterisks denote the degree of statistical significance (*: *p*-value < 0.05, **: *p*-value < 0.01, ***: *p*-value < 0.001).

## DISCUSSION

ROS such as H_2_O_2_, hydroxyl radicals, and carbonate radical anions are produced during oxidative stress and inflammation, contributing to aging and the development of many human pathologies including neurodegenerative disorders (74). Endogenous sources of ROS include aerobic respiration, inflammation, metabolism, and the immune response to infections (75). ROS induces damage to key cellular macromolecules, including DNA, proteins, and lipids, ultimately leading to genome instability and apoptotic cell death (7,59).Of those, DNA-protein cross-links are expected to be highly toxic due to their large molecular size and the ability to block normal DNA-protein interactions. If left unrepaired, DPCs can completely block vital biological functions such as replication, transcription, and repair, leading to cell death and mutations (1,2,18).

We have previously shown that ROS-derived DPCs can involve thymidine residues of DNA and tyrosine amino acid side chains of proteins to form Thy-Tyr conjugates **(C in Scheme 1)**. However, DPCs can form at other residues of cellular biomolecules (**Scheme 1**). Therefore, LC-MS analyses of Thy-Tyr adducts in treated cells potentially underestimate the total DPC load. While our previous studies have measured DPCs in mouse tissues, the identities of the participating proteins remained unknown, and the mechanisms of ROS-DPCs processing and removal in human cells were not well understood. We chose H_2_O_2_ as a model system to study ROS-mediated DNA damage in mammalian cells. H_2_O_2_ readily crosses biological membranes and, once inside the cells, can form more reactive ROS, such as hydroxyl radicals and carboxyl radicals anions in the presence of Fe^2+^ ions and carbonate salts (76).

In the present work, we investigated the formation and repair of H_2_O_2_-induced DNA-protein crosslinks (DPCs) in human cells. The K-SDS approach, similar to the one reported by Zhitkovich et al. (58), was used to estimate total DPC formation in cells in our study. Unlike LC-MS-based methods, this method is agnostic to DPC structure and does not allow for absolute quantification of DPC lesions. Calculated % DPC obtained by the K-SDS methodology is a relative measure of total DPC formation. We found this methodology to be extremely sensitive and useful for fast and efficient measurements of DPC formation and repair in cells. Several modifications to the original method were made, including the removal of RNA and quantification of DPC-containing DNA via Promega QuantFluor Assay kit **(Figure S1D)**. Using this methodology, we found that untreated cells contained quantifiable DPC amounts, but these levels increased in a concentration and time-dependent manner upon treatment with H_2_O_2_ (**Figure 1**). ROS-DPC formation was significantly prevented when HT1080 cells were pre-treated with a reducing agent (N-Ac-Cys) before H_2_O_2_ treatment **(Figure 1C)**. A similar effect was observed upon pre-treatment with 1 mM Vitamin C (Vit-C) **(Figure S5)**. Moreover, ROS-DPC levels gradually decreased after treatment, reaching background levels with an estimated half-life of 2 h (**Figure 1D**).

MS-based proteomics revealed that H_2_O_2_ treatment induced DPCs with many proteins of varying sizes, natures, and functions. Indeed, HT1080 cells treated with 200 µM H_2_O_2_ contained 105 DNA-crosslinked proteins varying in size, function, and cellular localization **(Figure 2C, Table S1)**. The most significant number of the identified proteins are of nuclear localization **(Figure S2A**) and are involved in DNA replication and transcription (e.g., IMPDH2, POLR2A, heterogeneous nuclear ribonucleoproteins), DNA repair (e.g., high mobility group proteins), and nucleosomal assembly (e.g., histones). A few ROS-induced DNA-crosslinked proteins, such as ubiquitin (UBB), heat shock protein 90 (HSP90), and tubulin alpha 1B (TUBA1B), have also been reported in our previous studies of endogenous DPC agents, notably methylglyoxal (5) and formaldehyde (manuscript in preparation).

Our study demonstrates that the proteolytic activities of both SPRTN metalloprotease and the ubiquitin-proteasomal system (UPS) could be important for removing ROS-DPCs from human cells. Indeed, reduced SPRTN protein expression sensitized cells to ROS-induced cytotoxicity **(Figure 3A)** and elevated the global levels of both endogenous and H_2_O_2_-derived DPCs, with more pronounced increase in DPC levels after inhibiting the ubiquitination pathway **(Figure 3B)**. This observation can be explained by the previous studies that reported ubiquitination-dependent SPRTN activity, including SPRTN recruitment to the DPC sites, both *in vitro* and in mammalian cells (77,78).

The ubiquitin-proteasomal system (UPS) is increasingly recognized as a parallel pathway through which DNA-trapped proteins are digested into peptides (28,50,66). However, similar studies have focused on the removal of DPCs induced by topoisomerase poisons, UV, and formaldehyde, leaving a gap in understanding whether UPS plays a role in the cellular clearance of ROS-DPCs. Co-treatment of HT1080 cells with H_2_O_2_ and the 26S proteasomal subunit inhibitor MG-132 resulted in a noteworthy increase in ROS-DPC levels **(Figure 3C)**, confirming a role of the proteasome in ROS-DPC removal.

Following proteolytic cleavage, the resulting DNA-peptide lesions must be repaired via canonical DNA repair pathways. Our results reveal a key role for nucleotide excision repair in removing ROS-induced DPCs in human cells. Indeed, genetic deficiency in the NER protein factors XPA, XPD, and XPF significantly increased DPC levels of DPCs in HT1080 cells **(Figure 4)**. We hypothesize that DNA-peptide crosslinks remaining on DNA following proteolytic processing by SPRTN or the proteasome are excised by NER machinery. This is consistent with previous studies of formaldehyde-induced DPCs (79,80). Furthermore, our results reveal an important role of XPF/ERCC1 in cellular response to ROS-DPCs. XPF forms a complex with ERCC1, and together they serve as a structure-specific endonuclease at the core of both transcription-coupled NER and global genome NER. While this protein complex was shown to participate in repairing other DNA damage besides DPCs, including interstrand cross-links (ICLs) and double-strand breaks (DSBs) (81,82), our K-SDS assays reveal a marked increase in DPC levels in cells deficient in XPF/ERCC1 **(Figure 4B)**, providing first direct evidence for XPF/ERCC1 involvement in ROS-DPC repair.

Collectively, our results are consistent with a model where H_2_O_2_-derived DPCs are detected by the replication and transcription machinery and undergo ubiquitination. Protein components of DPCs undergo both mono and polyubiquitination and are proteolytically digested by SPRTN metalloprotease or the proteasome. The resulting peptide lesions are removed by the GG-NER and TC-NER repair pathways **(Figure 3B, C&4B, C)**.

Finally, our work revealed the ability of the hydrolytically stable glutathione analog **ψ**-GSH (72) to prevent the formation of ROS-DPCs. Previously, supplementing glutathione (GSH) has been a tenable strategy to mitigate oxidative stress associated with several diseases, notably Alzheimer’s disease (AD) (83). A stable and cell-penetrating glutathione analog, **ψ**-GSH, showed promising *in vivo* therapeutic potential in AD mouse models (55). In the present work, *ψ*-GSH significantly reduced the global load of H_2_O_2_-derived DPCs and endogenous DPCs when H1080 cells were treated with higher concentrations of *ψ*-GSH **(Figure 5B)**. This effect can be attributed to GSH’s capacity to scavenge ROS and replenish endogenous GSH levels, a key cofactor of both glutathione peroxidase and glyoxalase (84).

In conclusion, this study demonstrates that exposure of human fibrosarcoma HT1080 cells to H_2_O_2_ induces cellular DPCs. MS-proteomics experiments identified 105 proteins involved in ROS-DPC formation. We confirmed that one of these proteins, HMGB2, can form DPCs *in vitro* when incubated with a synthetic DNA oligonucleotide in the presence of H_2_O_2_. Per our proposed model of cellular repair of ROS-DPCs **(Figure 6)**, when ROS-DPCs arise from exposure to ROS s **(step 1)**, they are detected by stalled replication and/or transcription machinery **(step 2)**. Ubiquitin ligases are recruited to the site of damage, leading to the ubiquitination of DNA-crosslinked proteins **(step 3)**. Next, SPRTN and the proteasome are recruited to degrade the ubiquitinated DPC proteins into smaller DNA-peptide crosslinks, which are accessible to the NER machinery **(step 4)**. During the NER pathway, the TFIIH and XPF-ERCC1 endonuclease complexes excise the DNA-peptide crosslinks followed by gap filling and ligation to produce intact DNA **(step 5)**. Collectively, our findings are consistent with several previous and emerging models of DPC repair reported elsewhere and offer important new details regarding ROS-DPC formation, processing and repair (7,18,36,41,78). Ongoing studies in our laboratory are optimizing analytical methods to assess the endogenous levels of specific ROS-DPC conjugates in several aging and AD tissue models and to determine the chemical structures of the resulting conjugates. Overall, these results provide a basis for studying the implications of ROS-derived DPCs in oxidative stress and related pathologies for future development of novel therapies.

**Figure 6:**
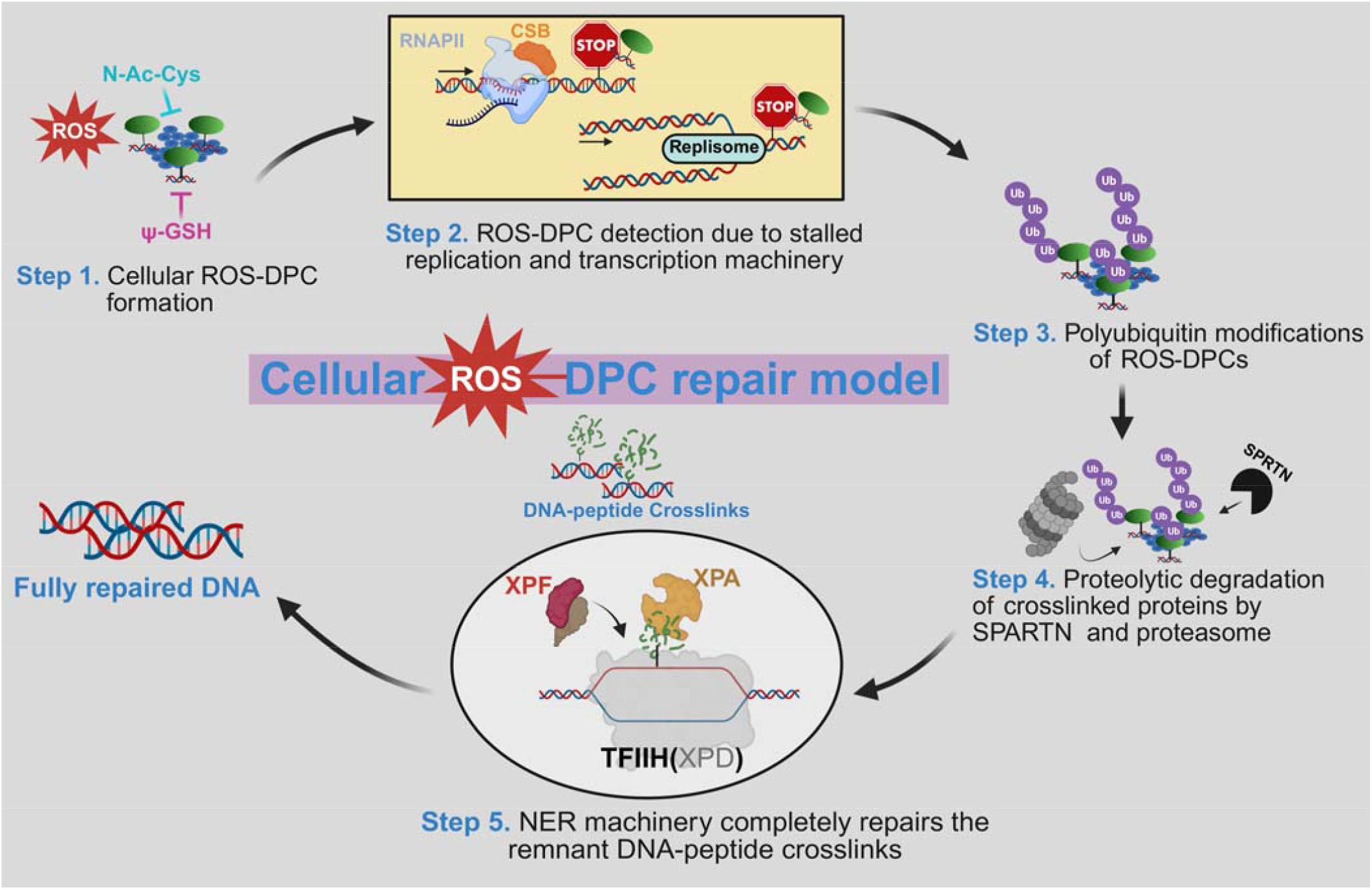
Proposed model of cellular repair of ROS-DPCs (figure created using *BioRender*)

## ACKNOWLEDGEMENTS

We thank Robert Carlson (University of Minnesota Masonic Cancer center) for improving the quality of the figures presented in this manuscript and other editorial help.

## AUTHOR CONTRIBUTIONS

Cesar I. Cyuzuzo (Conceptualization [supporting], Data curation [lead], Formal analysis [lead], Investigation [lead], Methodology [lead], Software [lead], Validation [equal], Visualization [lead], Writing—original draft [lead], writing—review & editing [lead], Monica Kruk (Investigation [supporting], Writing—review & editing [supporting]), Qi Zhang (Investigation [supporting], Writing—review & editing [supporting]), Duha Alshareef (Investigation [supporting], Writing—review & editing [supporting]), James Harmon (Investigation [supporting]), Yuichi J. Machida (Resources [supporting], Writing—review & editing [supporting]), Harrison W. VanKoten (Investigation [supporting]), Swati S. More (Resources [supporting], Writing—review & editing [supporting]), Colin Campbell (Conceptualization [supporting], Funding acquisition [equal], Resources [supporting], Supervision [supporting], Writing—review & editing [supporting]), and Natalia Y. Tretyakova (Conceptualization [lead], Data curation [equal], Formal Analysis [equal], Funding acquisition [lead], Methodology [equal], Project administration [lead], Resources [lead], Supervision [lead], Validation [equal], Visualization [equal], Writing—original draft [supporting], Writing—review & editing [equal].

## SUPPLEMENTARY DATA

Supplementary data are available at NAR online.

## DATA AVAILABILITY

Data underlying this article are available in the Zenodo public repository and can be accessed at DOI: 10.5281/zenodo.18175815

