## Supporting Information for "Living Cells Employ Ubiquitin-Proteasomal System and Nucleotide Excision Repair Pathways to Remove Reactive Oxygen Species-Induced DNA-Protein Crosslinks (ROS-DPCs)"

**Figure S1. Cross-validation of DNA quantitation assays**

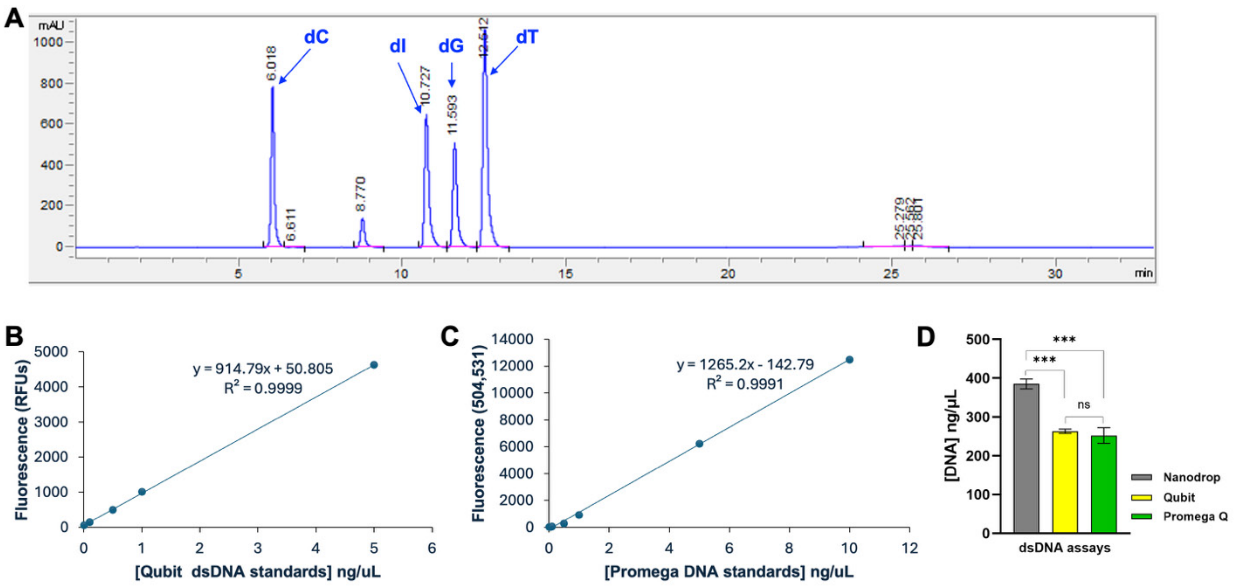


**(A)** Representative trace of dG analysis from $H_2O_2$-treated HT1080 cells. **(B)** DNA Standard curve for Qubit 3.0 fluorometry assay. **(C)** DNA Standard curve for Promega QuantiFluor (PQ) dsDNA system assay. **(D)** The estimated DNA amount using Nanodrop, Qubit, and PQ assay. DNA extracted from untreated HT1080 cells (n=3 biological replicates) via a K-SDS assay was measured using three DNA assays. DPC-associated DNA was estimated using Nanodrop (Grey), Qubit (Yellow), and Promega QuantFluor assay (Green). Data was analyzed via a one-way ANOVA and an unpaired *t*-test. Asterisks were used to denote the degree of statistical significance (∗: *p*-value < 0.05, ∗∗: *p*-value < 0.01, ∗∗∗: *p*-value < 0.001).

**Figure S2: Further analyses of mass spectrometry-based proteomics results for $H_2O_2$-DPCs**

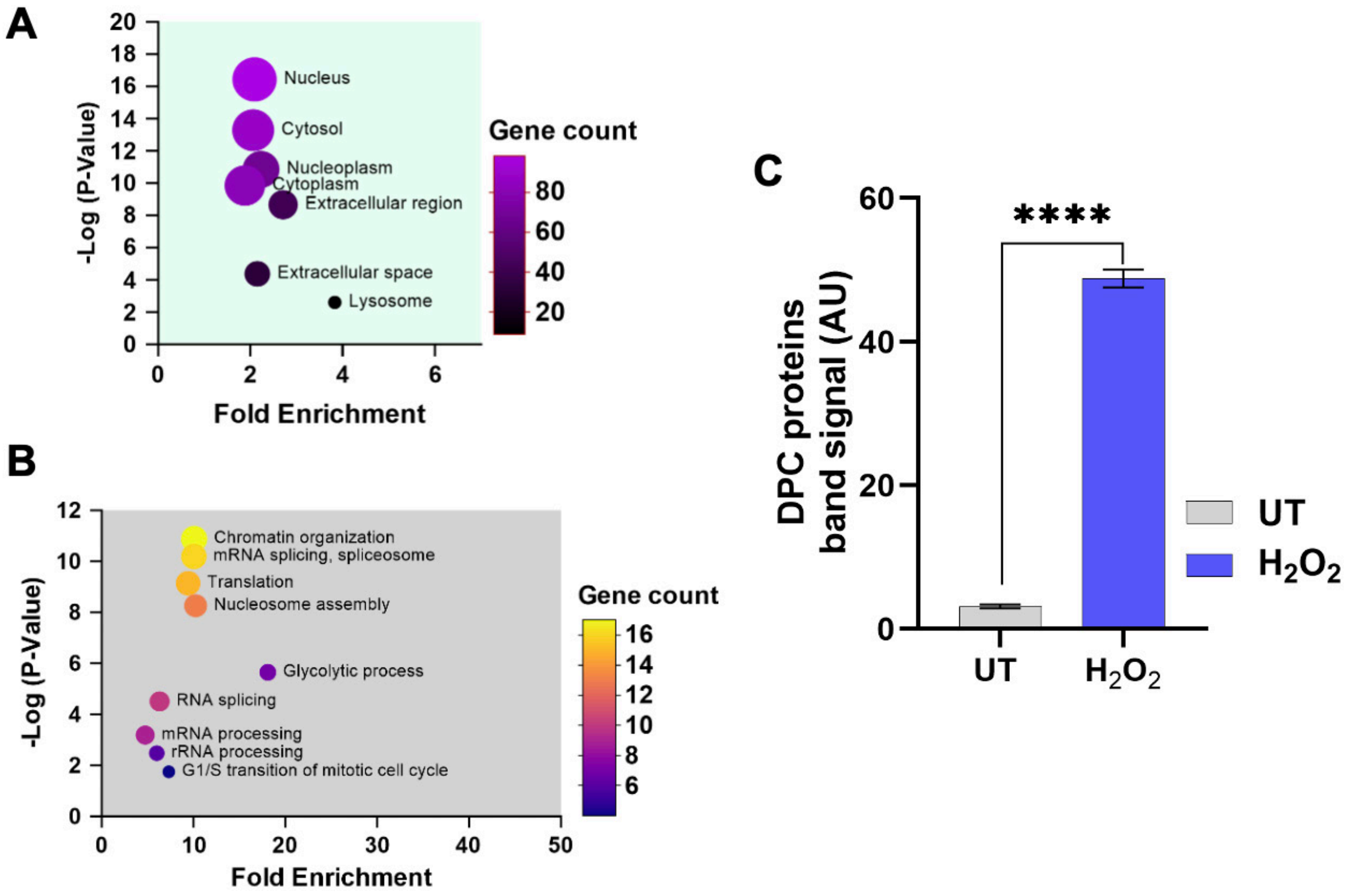


(**A**) Analysis was performed using the DAVID overrepresentation test, which produced Benjamini-Hochberg-adjusted *p*-values, gene counts, and fold enrichments for each enriched cellular compartment term. **(B)** A multivariate plot depicting the key biological processes of DNA-crosslinked proteins enriched by $H_2O_2$ treatment. Analysis was performed using the DAVID overrepresentation test, which produced Benjamini-Hochberg-adjusted *p*-values, gene counts, and fold enrichments for each enriched GO biological process term. (C) Densitometric analysis of the protein bands found in the 10-160 kDa molecular weight range in **Figure 2B**. Data were analyzed using one-way ANOVA and an unpaired *t*-test. Asterisks were used to denote the degree of statistical significance (*: *p*-value < 0.05, **: *p*-value < 0.01, ***: *p*-value < 0.001).

**Figure S3: Homologous recombination (HR) and autophagy pathways in cellular $H_2O_2$-DPC removal**

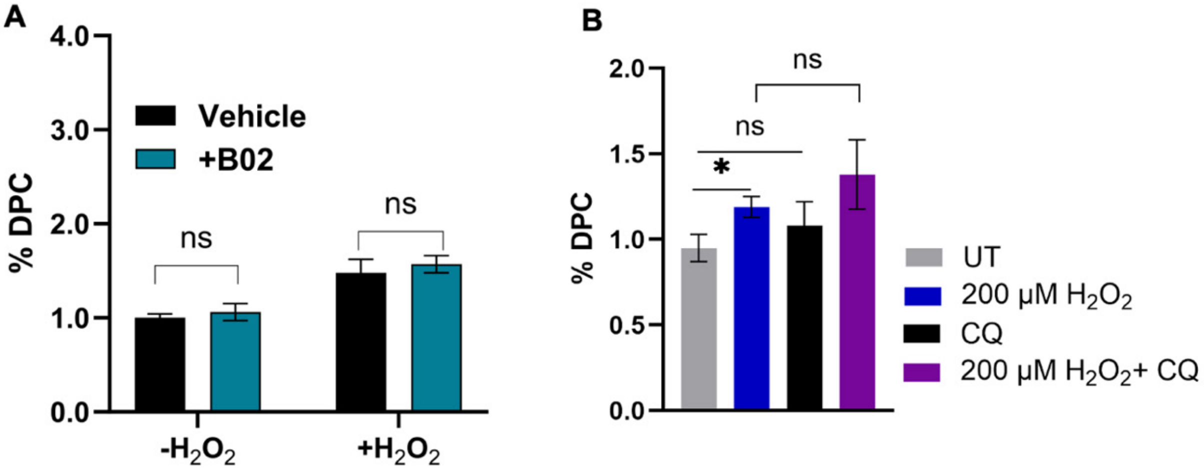


**(A)** HT1080 cells were co-treated with RAD51 recombinase inhibitor B02 and H2O2 (200 µM, 1 h, *n=3* biological replicates). DPC-associated DNA was extracted via K-SDS assay, and data were analyzed using one-way ANOVA and an unpaired *t*-test. Asterisks were used to denote the degree of statistical significance (∗: *p*-value < 0.05, ∗∗: *p*-value < 0.01, ∗∗∗: *p*-value < 0.001). **(B)** Effect of disrupting the autophagy pathway using chloroquine (CQ), an inhibitor that impairs autophagosome-lysosome fusion. In both **(A)** and **(B),** HT1080 cells were treated with CQ inhibitor in the presence or absence of $H_2O_2$ (200 µM, 1 h, *n=3* biological replicates). DPCs were isolated via a K-SDS assay, and data were analyzed using one-way ANOVA and an unpaired *t*-test. Asterisks were used to denote the degree of statistical significance (∗: *p*-value < 0.05, ∗∗: *p*-value < 0.01, ∗∗∗: *p*-value < 0.001).

**Figure S4: Western blot confirming the total loss of XPA in the HT080 cell line**

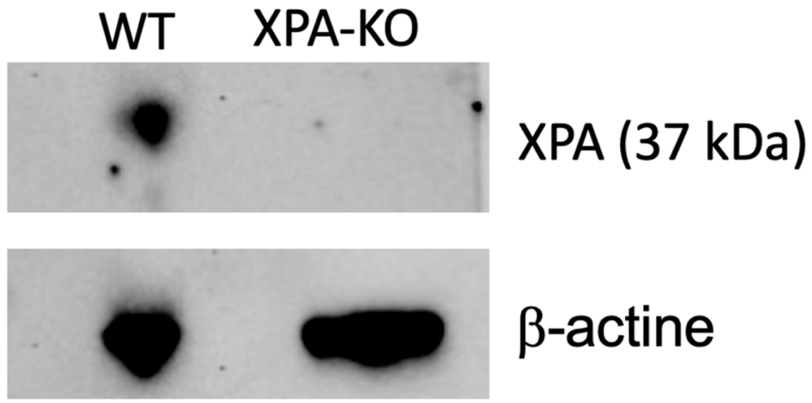


20 µg of nuclear protein extract from XPA-KO HT1080 cells was run on a 4-12% Bis-Tris SDS-PAGE gel. XPA and Beta-actin mouse monoclonal antibody (Santa Cruz Biotechnology) were primary antibodies. The goat anti-mouse HRP-conjugated secondary antibody (Invitrogen) was used, and the blot was developed using Pierce ECL Western Blotting Substrate (Thermo Scientific). The Image was taken using a Bio-Rad Imaging System.

**Figure S5: Higher amounts of vitamin C (L-ascorbic acid) scavenge $H_2O_2$**

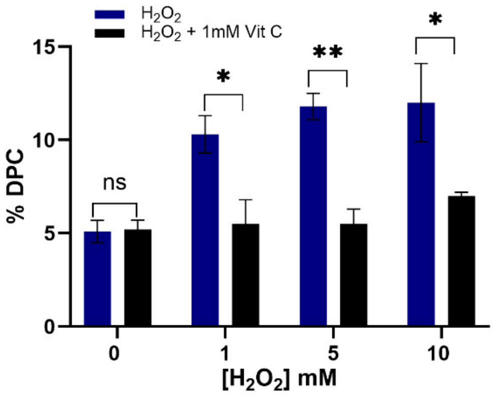


The vitamin C pre-treated HT080 cells were exposed to increasing amounts of $H_2O_2$ for 1 h, and after $H_2O_2$ quenching, a K-SDS assay was performed on collected pellets (*n=3* independent experiments). Data were analyzed using one-way ANOVA and an unpaired *t*-test. Asterisks were used to denote the degree of statistical significance (∗: *p*-value < 0.05, ∗∗: *p*-value < 0.01, ∗∗∗: *p*-value < 0.001).

**Table S1**: **DNA-crosslinked proteins enriched in the $H_2O_2$ treatment group**

The 3rd column depicts the unpaired t-test *p*-values. The 4th column shows the difference between the mean Log2-transformed LFQ intensities between the $H_2O_2$ and vehicle-treated groups. The 4th column values indicate the fold change difference in protein abundance. The data represent the results from three biological replicates used in each treatment condition.

| No. | Protein Name | Gene Name | Negative Log (*p*-value) | Difference ($H_2O_2$-Vehicle) |
|---|---|---|---|---|
| 1 | Neuroplastin | NPTN | 4.284 | 9.571 |
| 2 | Histone H3.1 | H3C12 | 1.975 | 9.361 |
| 3 | Phospholipase B-like 1 | PLBD1 | 0.790 | 9.129 |
| 4 | Lysosome-associated membrane glycoprotein 2 | LAMP2 | 3.804 | 8.615 |
| 5 | High mobility group protein B2 | HMGB2 | 3.105 | 8.586 |
| 6 | DNA-directed RNA polymerase II subunit RPB1 | POLR2A | 2.762 | 7.891 |
| 7 | Small ribosomal subunit protein uS5 | RPS2 | 2.471 | 7.652 |
| 8 | Inosine-5'-monophosphate dehydrogenase 2 | IMPDH2 | 5.301 | 7.479 |
| 9 | Small ribosomal subunit protein uS8 | RPS15A | 4.569 | 7.308 |
| 10 | GTP-binding nuclear protein Ran | RAN | 0.928 | 7.095 |
| 11 | Glucosamine-6-phosphate deaminase 1 | GNPDA1 | 0.625 | 6.957 |
| 12 | RNA-binding protein with serine-rich domain 1 | RNPS1 | 0.934 | 6.892 |
| 13 | High mobility group nucleosome-binding domain-containing protein 4 | HMGN4 | 0.903 | 6.811 |
| 14 | Large ribosomal subunit protein P1 | RPLP1 | 0.783 | 6.566 |
| 15 | T-complex protein 1 subunit eta | CCT7 | 0.933 | 6.303 |
| 16 | Polyadenylate-binding protein 1 | PABPC1 | 0.768 | 6.094 |
| 17 | Protein transport protein Sec61 subunit beta | SEC61B | 0.926 | 5.664 |
| 18 | High mobility group protein HMG-I/HMG-Y | HMGA1 | 2.038 | 3.625 |
| 19 | Branched-chain-amino acid aminotransferase | BCAT1 | 2.097 | 3.449 |
| 20 | RNA-binding motif protein, X chromosome | RBMX | 1.421 | 3.393 |
| 21 | High mobility group protein HMGI-C | HMGA2 | 2.549 | 3.356 |
| 22 | Beta-glucoronidase | GUSB | 1.213 | 3.218 |
| 23 | Trans-Golgi network integral membrane protein 2 | TGOLN2 | 0.795 | 3.018 |
| 24 | Serine/arginine repetitive matrix protein 1 | SRRM1 | 2.361 | 2.944 |
| 25 | Putative high mobility group protein B1-like 1 | HMGB1P1 | 2.667 | 2.922 |
| 26 | Protein arginine N-methyltransferase 1 | PRMT1 | 2.517 | 2.883 |
| 27 | Plastin-2 | LCP1 | 1.075 | 2.838 |
| 28 | Galectin-7 | LGALS7B | 1.967 | 2.837 |
| 29 | General transcription factor IIF subunit 1 | GTF2F1 | 2.835 | 2.836 |
| 30 | Protein disulfide-isomerase | P4HB | 2.384 | 2.821 |
| 31 | Dehydrogenase/reductase SDR family member 2, mitochondrial | DHRS2 | 0.362 | 2.806 |
| 32 | Nucleolin | NCL | 2.486 | 2.572 |
| 33 | Intestinal-type alkaline phosphatase | ALPI | 2.733 | 2.563 |
| 34 | Complement C3 | C3 | 0.599 | 2.497 |
| 35 | Serine/arginine-rich splicing factor 6 | SRSF6 | 1.616 | 2.427 |
| 36 | Endoplasmic reticulum chaperone BiP | HSPA5 | 1.466 | 2.391 |

| No. | Protein Name | Gene Name | Negative Log (*p*-value) | Difference ($H_2O_2$-Vehicle) |
|---|---|---|---|---|
| **37** | Alpha-N-acetylgalactosaminidase | NAGA | 1.872 | 2.359 |
| **38** | Acidic leucine-rich nuclear phosphoprotein 32 family member 2 | ANP32B | 2.272 | 2.335 |
| **39** | Transgelin-2 | TAGLN2 | 1.149 | 2.238 |
| **40** | Tubulin beta chain | TUBB | 1.644 | 2.224 |
| **41** | Immortalization up-regulated protein | IMUP | 0.924 | 2.217 |
| **42** | Desmoplakin | DSP | 1.365 | 2.198 |
| **43** | Basigin | BSG | 1.770 | 2.146 |
| **44** | Superoxide dismutase [Cu-Zn] | SOD1 | 1.137 | 2.136 |
| **45** | Eukaryotic translation initiation factor 4B | EIF4B | 1.371 | 2.07 |
| **46** | Bleomycin hydrolase | BLMH | 1.310 | 2.02 |
| **47** | Protein disulfide isomerase CRELD2 | CRELD2 | 0.304 | 2.005 |
| **48** | Cathepsin B | CTSB | 1.788 | 1.997 |
| **49** | ATP synthase subunit alpha, mitochondrial | ATP5F1A | 0.603 | 1.98 |
| **50** | 5'-3' exonuclease PLD3 | PLD3 | 1.420 | 1.895 |
| **51** | 14-3-3 protein epsilon | YWHAE | 0.293 | 1.889 |
| **52** | Nucleophosmin | NPM1 | 2.470 | 1.809 |
| **53** | Tubulin alpha-1B chain | TUBA1B | 1.719 | 1.738 |
| **54** | Non-histone chromosomal protein HMG-17 | HMGN2 | 0.766 | 1.707 |
| **55** | Alpha-enolase | ENO1 | 1.418 | 1.69 |
| **56** | Xaa-Pro dipeptidase | PEPD | 4.959 | 1.659 |
| **57** | Alkaline phosphatase, germ cell type | ALPG | 1.618 | 1.583 |
| **58** | Calreticulin | CALR | 0.851 | 1.563 |
| **59** | Small ribosomal subunit protein uS9 | RPS16 | 0.255 | 1.537 |
| **60** | Peroxiredoxin-4 | PRDX4 | 1.536 | 1.494 |
| **61** | Eukaryotic initiation factor 4A-I | EIF4A1 | 0.243 | 1.488 |
| **62** | Endoplasmin | HSP90B1 | 1.573 | 1.461 |
| **63** | Small ribosomal subunit protein uS2B | RPSA2 | 0.832 | 1.447 |
| **64** | Calnexin | CANX | 2.125 | 1.404 |
| **65** | Nucleoside diphosphate kinase B | NME2 | 1.533 | 1.402 |
| **66** | Dolichyl-disphosphooligosaccharide-protein glycosyltransferase 48 kDa subunit | DDOST | 0.423 | 1.362 |
| **67** | Protein CutA | CUTA | 0.650 | 1.331 |
| **68** | C(dfdD44 anginer) | CD44 | 2.464 | 1.315 |
| **69** | Glyceraldehyde-3'-phosphate dehydrogenase | GAPDH | 1.079 | 1.298 |
| **70** | Rab GDP dissociation inhibitor alpha | GDI2 | 0.359 | 1.222 |
| **71** | Elongation factor 2 | EEF2 | 2.460 | 1.202 |
| **72** | Small ribosomal subunit protein eS6 | RPS6 | 0.424 | 1.18 |
| **73** | Lysosome membrane protein 2 | SCARB2 | 0.466 | 1.149 |
| **74** | Integrin beta-1 | ITGB1 | 1.501 | 1.13 |
| **75** | T-complex protein 1 subunit | CCT8 | 0.823 | 1.103 |
| **76** | Heterogeneous nuclear ribonucleoprotein K | HNRNPK | 1.874 | 1.1 |
| **77** | Serine/arginine repetitive matrix protein 2 | SRRM2 | 1.338 | 1.091 |
| **78** | Triosephosphate isomerase | TPI1 | 0.298 | 1.032 |

| No. | Protein Name | Gene Name | Negative Log (*p*-value) | Difference ($H_2O_2$-Vehicle) |
|---|---|---|---|---|
| **79** | 14-3-3 protein zeta/delta | YWHAZ | 0.387 | 0.999 |
| **80** | Lysosome-associated membrane glycoprotein 1 | LAMP1 | 0.327 | 0.935 |
| **81** | Actin, cytoplasmic 1 | ACTB | 1.954 | 0.848 |
| **82** | Keratin, type I cytoskeletal 9 | KRT9 | 0.517 | 0.846 |
| **83** | Peroxiredoxin-1 | PRDX1 | 0.316 | 0.837 |
| **84** | Large ribosomal subunit protein uL5 | RPL11 | 0.297 | 0.817 |
| **85** | Small ribosomal subunit protein uS4 | RPS9 | 0.269 | 0.803 |
| **86** | Pyruvate kinase PKM | PKM | 1.511 | 0.776 |
| **87** | Lysine-tRNA ligase | KARS1 | 0.140 | 0.774 |
| **88** | Heat shock protein HSP 90-beta | HSP90AB1 | 2.223 | 0.756 |
| **89** | Tubulin alpha-1A chain | TUBA1A | 2.496 | 0.743 |
| **90** | Profilin-1 | PFN1 | 1.673 | 0.742 |
| **91** | Small ribosomal subunit protein uS3 | RPS3 | 0.205 | 0.652 |
| **92** | Elongation factor 1-alpha 1 | EEF1A1 | 0.860 | 0.65 |
| **93** | Nucleolar and coiled-body phosphoprotein 1 | NOLC1 | 0.259 | 0.622 |
| **94** | Vimentin | VIM | 1.327 | 0.588 |
| **95** | Thioredoxin reductase 1, cytoplasmic | TXNRD1 | 0.098 | 0.563 |
| **96** | Annexin A2 | ANXA2 | 0.944 | 0.537 |
| **97** | Translocation protein SEC62 | SEC62 | 0.235 | 0.504 |
| **98** | Heterogeneous nuclear ribonucleoprotein U | HNRNPU | 0.132 | 0.486 |
| **99** | Histone H2B type 1-K | H2BC12 | 0.515 | 0.45 |
| **100** | Amino acid transporter | SLC1A5 | 0.082 | 0.406 |
| **101** | L-lactate dehydrogenase A chain | LDHA | 0.783 | 0.397 |
| **102** | Polyubiquitin-B | UBB | 1.361 | 0.392 |
| **103** | Albumin | ALB | 0.246 | 0.313 |
| **104** | Aspartyl/asparaginyl beta-hydroxylase | ASPH | 0.138 | 0.206 |
| **105** | keratin, type II cytoskeletal 2 epidermal | KRT2 | 0.056 | 0.199 |
